# Neoadjuvant systemic oncolytic vesicular stomatitis virus is safe and may enhance long-term survivorship in dogs with naturally occurring osteosarcoma

**DOI:** 10.1101/2023.04.16.533664

**Authors:** Kelly M. Makielski, Aaron L. Sarver, Michael S. Henson, Kathleen M. Stuebner, Antonella Borgatti, Lukkana Suksanpaisan, Caitlin Preusser, Alexandru-Flaviu Tabaran, Ingrid Cornax, M. Gerard O’Sullivan, Andrea Chehadeh, Donna Groschen, Kelly Bergsrud, Sara Pracht, Amber Winter, Lauren J. Mills, Marc D. Schwabenlander, Melissa Wolfe, Michael A. Farrar, Gary R. Cutter, Joseph S. Koopmeiners, Stephen J. Russell, Jaime F. Modiano, Shruthi Naik

## Abstract

Osteosarcoma is a devastating bone cancer that disproportionally afflicts children, adolescents, and young adults. Standard therapy includes surgical tumor resection combined with multiagent chemotherapy, but many patients still suffer from metastatic disease progression. Neoadjuvant systemic oncolytic virus (OV) therapy has the potential to improve clinical outcomes by targeting primary and metastatic tumor sites and inducing durable antitumor immune responses. Here we described the first evaluation of neoadjuvant systemic therapy with a clinical-stage recombinant oncolytic Vesicular stomatitis virus (VSV), VSV-IFNβ-NIS, in naturally occurring cancer, specifically appendicular osteosarcoma in companion dogs. Canine osteosarcoma has a similar natural disease history as its human counterpart. VSV-IFNβ-NIS was administered prior to standard of care surgical resection, permitting microscopic and genomic analysis of tumors. Treatment was well-tolerated and a “tail” of long-term survivors (∼35%) was apparent in the VSV-treated group, a greater proportion than observed in two contemporary control cohorts. An increase in tumor inflammation was observed in VSV-treated tumors and RNAseq analysis showed that all the long-term responders had increased expression of a T-cell anchored immune gene cluster. We conclude that neoadjuvant VSV-IFNβ-NIS is safe and may increase long-term survivorship in dogs with naturally occurring osteosarcoma, particularly those that exhibit pre-existing antitumor immunity.

## Introduction

Osteosarcoma is a devastating bone cancer that primarily affects children, adolescents, and young adults, occurring most commonly in the long bones of the limbs.^1^ Standard treatment for patients diagnosed with osteosarcoma is definitive tumor resection with limb-sparing surgery elected where possible, combined with multi-agent chemotherapy administered in the neoadjuvant and adjuvant settings. These protocols were developed over 40 years ago, achieving a 5-year survival rate of approximately 60%.^1,2^ Many patients have inoperable metastases (commonly in the lung) and mortality is often due to progression of pre-existing or new sites of metastatic disease. Patients who are refractory to frontline treatment or are diagnosed with metastatic osteosarcoma have a very poor prognosis.^3^ Treatment modifications tested to date have failed to have a major impact on clinical outcomes. Thus, there is an unmet clinical need for treatments that improve outcomes for osteosarcoma patients. Development of new treatments is hampered by the rarity of this malignancy. Naturally occurring canine cancer represents a clinically relevant model that can recapitulate the spontaneous development and heterogeneity present in human cancer.^4^ Osteosarcoma is a common malignancy in dogs with similar clinical presentation and natural history as in humans. Osteosarcoma treatment in dogs includes surgical amputation of the affected limb and adjuvant chemotherapy.^5^ Also as in humans, despite removal of the primary tumor, most dogs develop pulmonary metastases and succumb to the disease; however, in dogs this typically occurs in less than 1 year.^6^

Oncolytic viruses (OVs) are engineered to selectively kill tumor cells and recruit immune cells to tumor sites.^7^ Vesicular stomatitis virus (VSV) expressing interferon-beta (IFNβ) and the sodium iodide symporter (NIS), referred to henceforth as VSV-IFNβ-NIS, is a clinical-stage recombinant OV that is currently being tested as a systemic therapy for patients with advanced solid tumors and hematologic malignancies both as a monotherapy and in combination with immune checkpoint inhibitors.^8,9^ Preclinical studies showed that systemically administered VSV infected and amplified selectively in tumor cells inducing tumor cell death, antitumor immune responses, and durable tumor remission in murine tumor models.^10–13^ Antitumor activity of systemic VSV therapy was also demonstrated in naturally occurring cancer in dogs. Single dose systemic VSV-IFNβ-NIS therapy was well tolerated with clinical remissions of disseminated lesions observed in two dogs with lymphoma and disease stabilization in a dog with metastatic osteosarcoma.^14^ These early clinical responses were also notable because most of the dogs had received prior treatments (mainly chemotherapy) and were enrolled after diagnosis with advanced refractory or relapsed disease. Our goal in this study was to evaluate the use of systemic VSV-IFNβ-NIS therapy in the neoadjuvant setting in dogs with newly diagnosed, localized osteosarcoma.

Systemic OV therapy administered in the neoadjuvant setting has the potential to target and kill malignant cells in primary and metastatic tumor sites, inducing inflammation in intact tumors to generate antitumor immune responses that persist after surgical interventions to improve clinical outcomes following standard of care treatment.^15^ Due to the prevalence of metastatic pulmonary recurrence in canine (and human) osteosarcoma, we hypothesized that systemic VSV therapy administered in the neoadjuvant setting has the potential to improve clinical outcomes following standard therapy for osteosarcoma. We launched the VIGOR study (VSV Immunotherapy and Genomics of Osteosarcoma Research) to test the safety, efficacy, and immunomodulatory effects of neoadjuvant intravenous VSV-IFNβ-NIS therapy in dogs with localized appendicular osteosarcoma.

## Results

### Oncolytic VSV kills canine osteosarcoma cells *in vitro*

Type I interferons, including IFNβ, activates antiviral innate immune responses in normal cells. Oncolytic VSV expressing IFNβ has been shown to selectively replicate within and kill a variety of tumor cells due to impaired innate immune responses.^16–18^ We previously generated and characterized a recombinant VSV expressing canine IFNβ and NIS (VSV-cIFNβ-NIS) for use in veterinary clinical trials in dogs.^14^ We evaluated the replication and oncolytic activity of recombinant VSV vectors, VSV-GFP, VSV-hIFNβ-NIS (VSV expressing human IFNβ), and VSV-cIFNβ-NIS in canine osteosarcoma cell lines (OSCA-78, OSCA-08, and OSCA-40) *in vitro*. Evaluation of VSV expressing canine and human IFNβ in canine osteosarcoma cells permits comparison of the effect of VSV expressing active canine IFNβ in canine cells. The VSV vectors replicated in all three canine osteosarcoma cell lines, with VSV-IFNβ-NIS vectors having ∼1-log lower maximal titer compared to VSV-GFP. VSV-IFNβ-NIS (expressing both human and canine IFNβ) replication resulted in tumor cell killing with similar potency as VSV-GFP resulting in <50% viability ∼48 hours post infection (Figure 1). These data confirm that canine osteosarcoma cell lines are susceptible to VSV infection and oncolysis and expression of canine IFNβ does not attenuate VSV replication and oncolysis in canine osteosarcoma cell lines.

**Figure 1.**
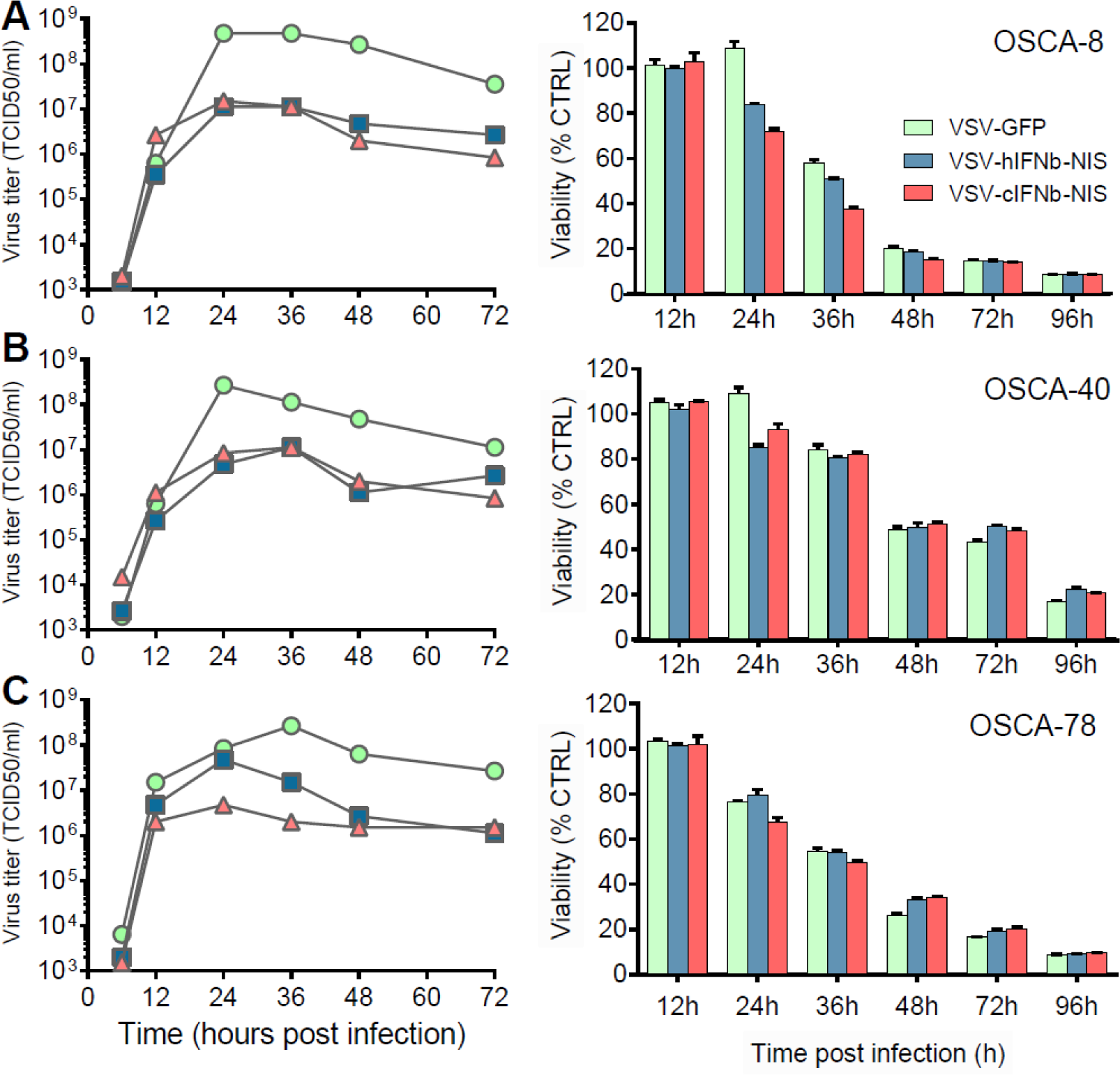
Vesicular stomatitis virus replicates in and kills three established canine osteosarcoma cell lines. Virus titers and cell viability following infection of canine osteosarcoma cell lines: (A) OSCA-8, (B) OSCA-40, and (C) OSCA-78 with recombinant VSVs: VSV-GFP (green), VSV-hIFNβ-NIS (blue), or VSV-cIFNβ-NIS (pink) at an MOI of 0.03.

### Screening and enrollment to the VIGOR study

The VIGOR (**<u>V</u>**SV **<u>I</u>**mmunotherapy and **<u>G</u>**enomics for **<u>O</u>**steosarcoma **<u>R</u>**esearch) study outline is shown in Figure 2. A total of 41 dogs were fully screened for enrollment in the VIGOR study of the 144 inquiries received (Supplemental Figure S1). Of the dogs screened, 13 were excluded. Twelve of these did not meet eligibility criteria due to documentation of metastatic disease on screening diagnostics (n=4), clinical suspicion or diagnosis of a different cancer (n=4), evidence of a pathologic fracture (n=3), or concurrent metastatic disease and pathologic fracture (n=1). One screened dog was not enrolled because the owner declined to participate. The study enrolled a total of 28 dogs over a period of 30 months (June 2016 to January 2019) with demographic characteristics of the dogs enrolled in the VIGOR study, as well as the contemporary control comparison populations, shown in Table 1. All dogs had an initial diagnosis of sarcoma of an appendicular bone based on a pre-treatment bone biopsy. This was a fixed-dose study with the first 15 dogs receiving one dose of 1×10^9^ TCID_50_/kg neoadjuvant VSV-cIFNβ-NIS under an open label design. After the safety of systemic neoadjuvant VSV-cIFNβ-NIS was documented, the next 13 dogs were randomized in a double-blinded fashion to receive a single dose of either intravenous VSV-cIFNβ-NIS (n=7) or placebo (PBS, n=6). All dogs were subsequently treated with standard of care and underwent amputation of the affected limb and removal of the associated lymph node(s) 10 days after receiving VSV-cIFNβ-NIS or placebo. Faxitron imaging was performed on the amputated limb to guide sample collection (Supplemental Figure S2). Pathological analysis of resected tumors confirmed primary osteosarcoma in 26 dogs, but the diagnoses in two dogs were subsequently revised: one (treated with VSV in the open label portion of the study) was diagnosed with intramedullary hemangiosarcoma at amputation, and the other (randomized to receive placebo) was diagnosed with intramedullary rhabdomyosarcoma at necropsy.

**Figure 2.**
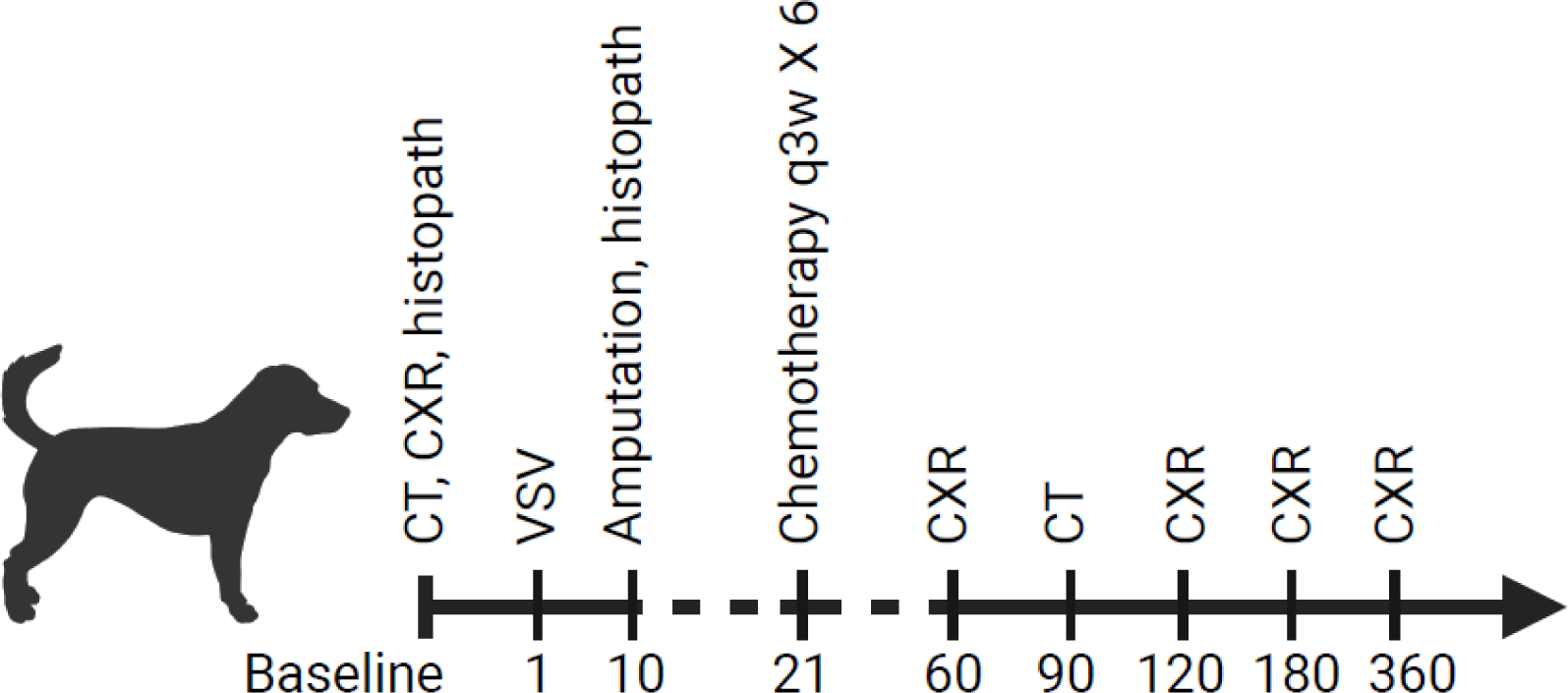
Design schematic of the VIGOR study. Clinical procedures following enrollment of dogs with appendicular osteosarcoma are shown. This includes diagnostic imaging to evaluate for pulmonary metastases with either computed tomography (CT) or thoracic radiographs (CXR); pre-treatment biopsy; VSV (or placebo) treatment (study day 1); and standard of care amputation and chemotherapy (consisting of intravenous carboplatin every 3 weeks for 6 doses) that was started on study day 21.

**Table 1.**
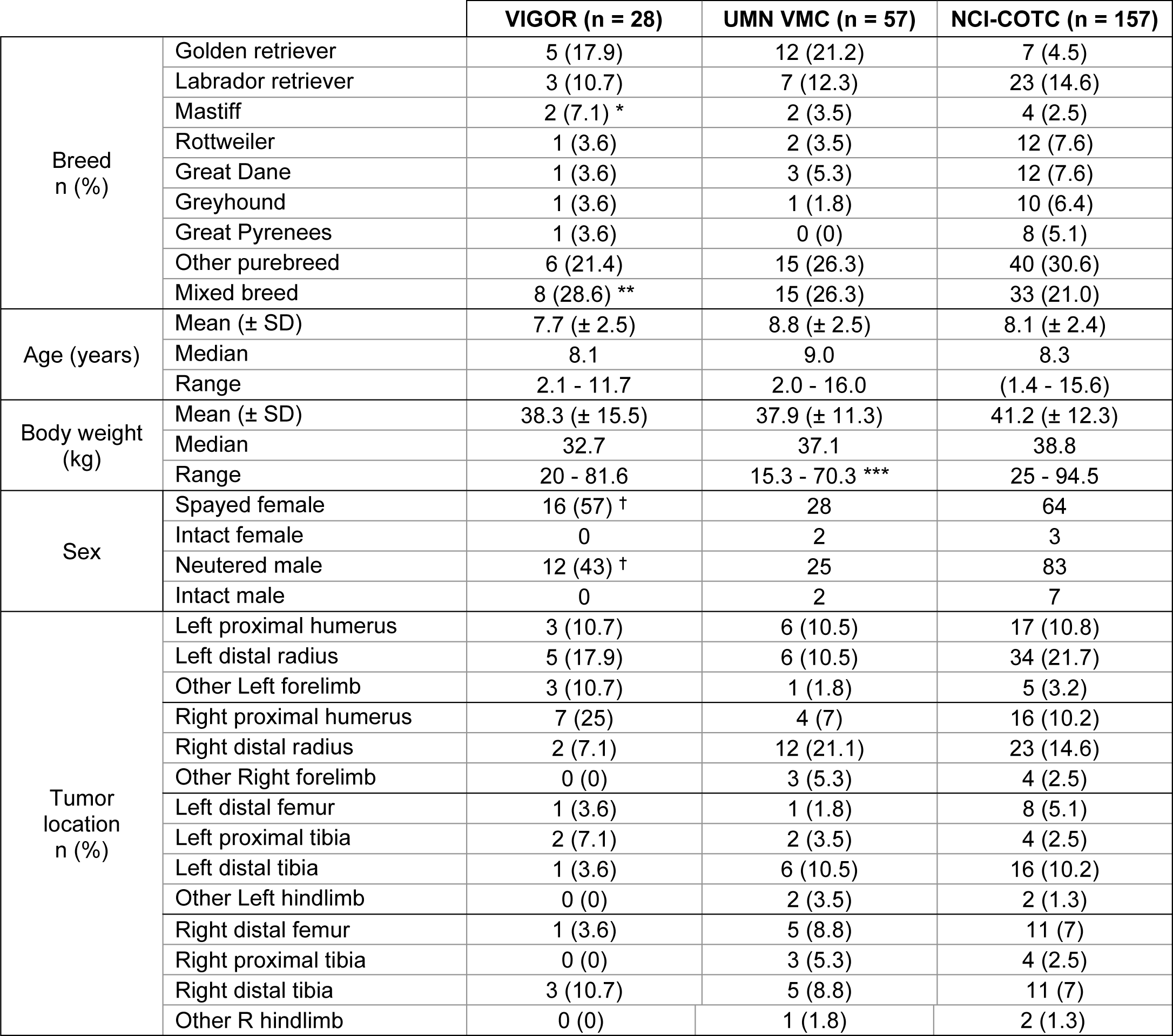
The clinical characteristics of the dogs enrolled in the VIGOR study are similar to contemporary comparison cohorts with appendicular osteosarcoma. Dogs with osteosarcoma tumors in the appendicular skeleton were enrolled in VIGOR. The demographic characteristics of the dogs enrolled in the VIGOR cohort were generally large breed middle-aged dogs. VIGOR: test population; UMN VMC: comparison cohort at same site (University of Minnesota Veterinary Medical Center); COTC: national multicenter comparison cohort overseen by the Comparative Oncology Network at the NCI. VIGOR, UMN, and COTC were all treated with the same standard of care chemotherapy. All cases were treated with amputation. * One Mastiff in VIGOR was diagnosed with intramedullary rhabdomyosarcoma. ** One mixed breed enrolled in VIGOR was diagnosed with intramedullary hemangiosarcoma. *** Body weight was not available for 5 of the dogs in the UMN comparison group. ^†^ All dogs were neutered and weighed > 20kg as conditions of enrollment in VIGOR.

|  |  | VIGOR (n = 28) | UMN VMC (n = 57) | NCI-COTC (n = 157) |
| --- | --- | --- | --- | --- |
| Breed<br>n (%) | Golden retriever | 5 (17.9) | 12 (21.2) | 7 (4.5) |
|  | Labrador retriever | 3 (10.7) | 7 (12.3) | 23 (14.6) |
|  | Mastiff | 2 (7.1) * | 2 (3.5) | 4 (2.5) |
|  | Rottweiler | 1 (3.6) | 2 (3.5) | 12 (7.6) |
|  | Great Dane | 1 (3.6) | 3 (5.3) | 12 (7.6) |
|  | Greyhound | 1 (3.6) | 1 (1.8) | 10 (6.4) |
|  | Great Pyrenees | 1 (3.6) | 0 (0) | 8 (5.1) |
|  | Other purebreed | 6 (21.4) | 15 (26.3) | 40 (30.6) |
|  | Mixed breed | 8 (28.6) ** | 15 (26.3) | 33 (21.0) |
| Age (years) | Mean (± SD) | 7.7 (± 2.5) | 8.8 (± 2.5) | 8.1 (± 2.4) |
|  | Median | 8.1 | 9.0 | 8.3 |
|  | Range | 2.1 - 11.7 | 2.0 - 16.0 | (1.4 - 15.6) |
| Body weight<br>(kg) | Mean (± SD) | 38.3 (± 15.5) | 37.9 (± 11.3) | 41.2 (± 12.3) |
|  | Median | 32.7 | 37.1 | 38.8 |
|  | Range | 20 - 81.6 | 15.3 - 70.3 *** | 25 - 94.5 |
| Sex | Spayed female | 16 (57) <sup>†</sup> | 28 | 64 |
|  | Intact female | 0 | 2 | 3 |
|  | Neutered male | 12 (43) <sup>†</sup> | 25 | 83 |
|  | Intact male | 0 | 2 | 7 |
| Tumor<br>location<br>n (%) | Left proximal humerus | 3 (10.7) | 6 (10.5) | 17 (10.8) |
|  | Left distal radius | 5 (17.9) | 6 (10.5) | 34 (21.7) |
|  | Other Left forelimb | 3 (10.7) | 1 (1.8) | 5 (3.2) |
|  | Right proximal humerus | 7 (25) | 4 (7) | 16 (10.2) |
|  | Right distal radius | 2 (7.1) | 12 (21.1) | 23 (14.6) |
|  | Other Right forelimb | 0 (0) | 3 (5.3) | 4 (2.5) |
|  | Left distal femur | 1 (3.6) | 1 (1.8) | 8 (5.1) |
|  | Left proximal tibia | 2 (7.1) | 2 (3.5) | 4 (2.5) |
|  | Left distal tibia | 1 (3.6) | 6 (10.5) | 16 (10.2) |
|  | Other Left hindlimb | 0 (0) | 2 (3.5) | 2 (1.3) |
|  | Right distal femur | 1 (3.6) | 5 (8.8) | 11 (7) |
|  | Right proximal tibia | 0 (0) | 3 (5.3) | 4 (2.5) |
|  | Right distal tibia | 3 (10.7) | 5 (8.8) | 11 (7) |
|  | Other R hindlimb | 0 (0) | 1 (1.8) | 2 (1.3) |

### VSV-cIFNB-NIS treatment is safe and is associated with prolonged survival in a subset of dogs with non-metastatic appendicular osteosarcoma

No clinically significant laboratory abnormalities were observed following VSV administration. About half the treated dogs developed a transient fever (> 1.5°C increase in body temperature) that resolved without intervention within 72 hours post infusion (not shown). Transient hepatotoxicity and lymphopenia are commonly observed following intravenous VSV infusion.^8,14^ As expected, mild lymphopenia was observed in some dogs following VSV infusion that resolved spontaneously (Supplemental Figure 3). There were no severe adverse events (AEs) directly attributable to VSV-cIFNβ-NIS. Three dogs that were treated with VSV (two open-label and one randomized to receive VSV) had reportable AEs that were considered independent from the expected side effects of chemotherapy (Supplemental Table S1), including one case each of pneumonia, facial nerve paralysis, and hypovolemic shock after amputation. The latter were post-surgical complications that resulted in death of the dog and likely not related to VSV-cIFNβ-NIS administration.

Clinical efficacy endpoints evaluated included event free survival (EFS) and overall survival. Overall survival for dogs in the VIGOR study roughly segregated into 3 populations with a “tail” of 7 dogs that were “long-term” survivors. The group of long-term survivors includes 4 dogs that are still alive at the time of this report including three dogs with osteosarcoma (two treated with VSV and one randomized to receive placebo) and one with hemangiosarcoma of bone (treated with VSV). Three dogs were excluded from the survival analysis: two dogs that eventually received a histopathologic diagnosis other than osteosarcoma (one rhabdomyosarcoma of bone (placebo-treated) and one hemangiosarcoma of bone (VSV-treated)) and one dog that died immediately after surgery due to hypovolemic shock (VSV-treated), yielding a cohort of 20 evaluable VSV-cIFNβ-NIS treated dogs. The clinical information, treatment categories, and survival data for each individual dog enrolled in VIGOR are shown in Supplementary Table S2. EFS and overall survival for these evaluable dogs were compared to two contemporary control cohorts of dogs with appendicular osteosarcoma with no evidence of metastasis. The first cohort (n = 57) included dogs seen at the University of Minnesota (UMN) VMC between July 2011 and July 2018, where there was intent to treat with standard-of-care surgery and adjuvant carboplatin chemotherapy, and that had a successful limb amputation and completed at least one cycle of adjuvant chemotherapy. The second control cohort was from a study recently published by the National Cancer Institute’s Comparative Oncology Trial Consortium (NCI-COTC) and included 157 dogs between November 2015 and February 2018 enrolled at 18 sites around the United States that were treated with standard-of-care surgery and adjuvant carboplatin chemotherapy.^19^ The NCI-COTC national multicenter comparison cohort was made available as a control dataset for evaluation of novel neoadjuvant or adjuvant treatments for osteosarcoma. Comparison to control cohorts showed that neoadjuvant VSV-cIFNβ-NIS therapy did not markedly improve EFS or overall survival (Figure 3A), but importantly did not worsen survival outcomes. Based on the NCI-COTC control cohort, “long-term” survivorship was defined as overall survival that exceeded the 75th percentile value of the NCI-COTC cohort (479 days). The UMN VMC control cohort had, as expected, 26% of dogs that exceeded an overall survival of 479 days. The VIGOR cohort had a higher-than-expected proportion of long-term survivorship with 35% of dogs exceeding an overall survival of 479 days (Figure 3B), though the sample size lacked the power to establish that this difference was not due to random chance with a high level of confidence.

**Figure 3.**
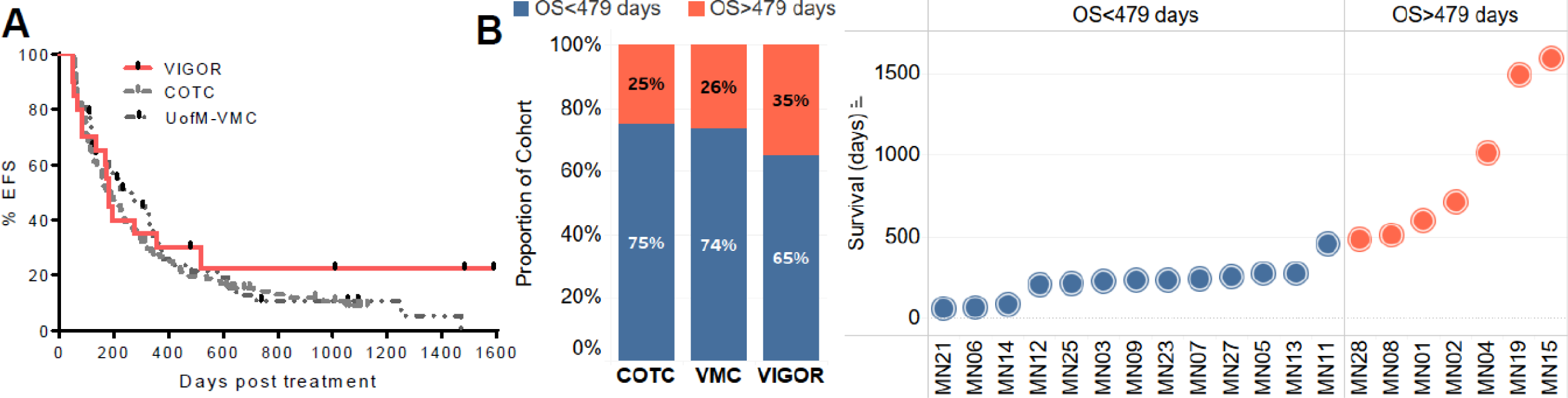
Intravenous VSV therapy is safe, and a subset of VSV-treated dogs with osteosarcoma are long term responders. (A) Kaplan-Meier survival curve with event free survival (EFS) of dogs treated with systemic VSV in the VIGOR study, compared to two contemporary control cohorts: NCI-COTC and UMN VMC cohorts. (B) Long-term response was defined as overall survival greater then 75^th^ percentile of survival from the COTC cohort (479 days). Similar proportion (26%) of dogs from the UMN VMC cohort were long-term responders. 35% of dogs from the VSV-treated dogs from the VIGOR cohort were long term responders (overall survival > 479 days).

### Histopathological assessment of osteosarcoma tumors shows an increase in tumor inflammation following VSV treatment

Pre-treatment tumor biopsies and post-treatment amputation-resected tumor samples were scored based on tumor necrosis, inflammation, and fibrosis (supplemental table S3). Tumor histopathologic evaluation showed areas of moderate to severe ischemic necrosis, as is typically observed in rapidly growing osteosarcoma lesions, in both VSV-cIFNβ-NIS-treated and placebo-treated cases; however, the tumors of 10 of 22 VSV-cIFNβ-NIS-treated dogs had distinctive areas of micronecrosis that were not consistently present in placebo-treated tumors (Supplemental Figure S4). Inflammatory infiltrates were scored in pre-treatment and post-treatment tumor specimens. Higher tumor infiltration score (TIS) was observed in post-treatment tumor specimens from VSV treated dogs compared to pre-treatment tumor biopsies, while similar increases were not observed in tumor specimens from placebo treated dogs (Figure 4A). Matched pre-vs post-treatment tumor specimens were available for assessment of intratumoral inflammatory infiltrate for half of the enrolled dogs showing a significant increase in TIS in post-treatment tumors compared to baseline (Figure 4B, P=0.0027). These results are encouraging but must be interpreted with caution given the small cohort size and the small number of dogs enrolled in the placebo group.

**Figure 4.**
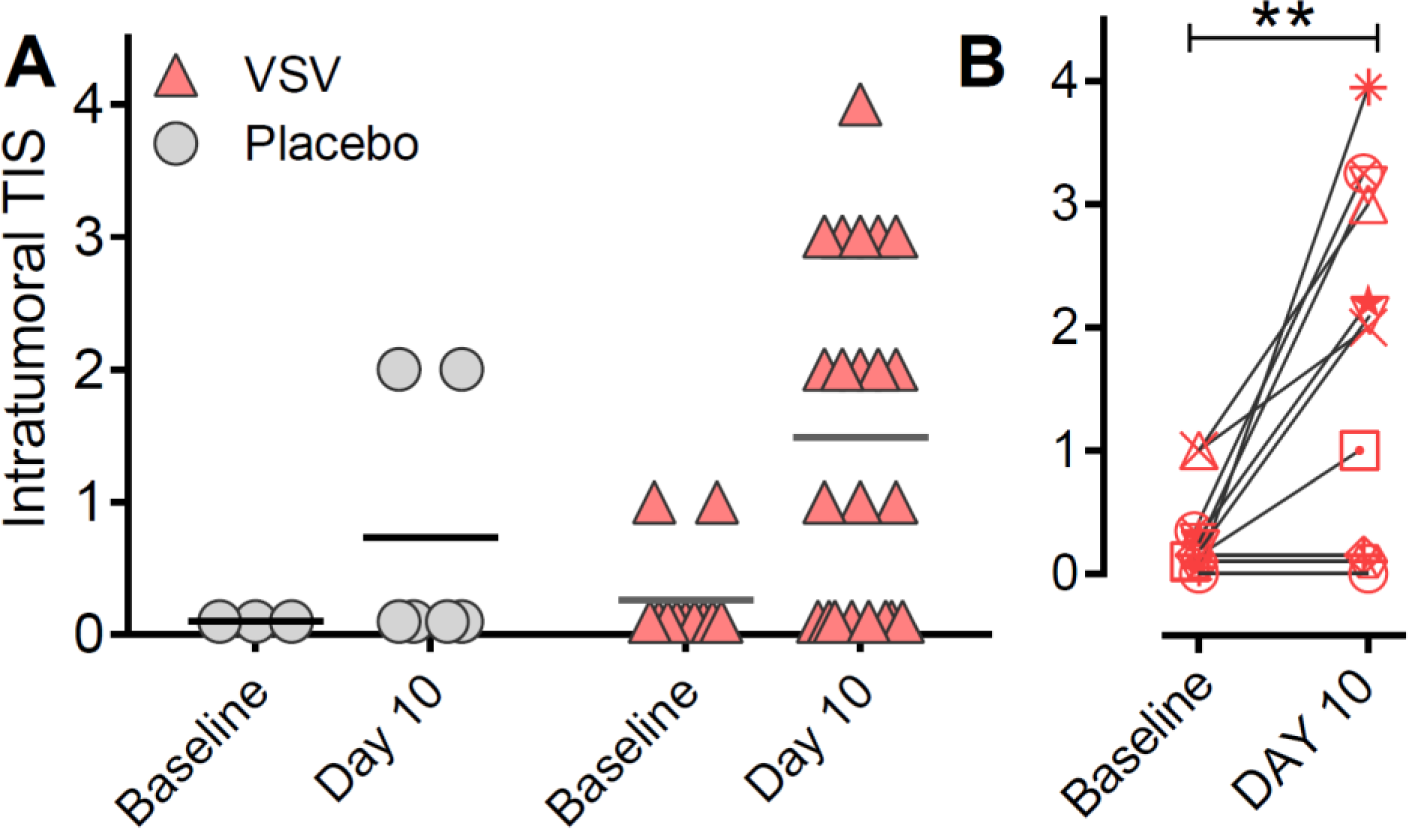
Increased tumor inflammation following VSV treatment in resected osteosarcoma tissues. Inflammatory infiltrates were scored by a pathologist in baseline pre-treatment tumor biopsies and post-treatment amputation-resected tumor samples (collected on study day 10). (A) Comparison of Tumor Inflammation Score (TIS) showed higher TIS in post-treatment samples from VSV-treated dogs. (B) Where matched paired pre- and post-treatment tumor samples were available for assessment, we observed a significant increase in TIS in tissues from VSV-treated dogs between baseline and 10 days post treatment (P=0.0027, paired t-test).

Regional lymph nodes were evaluated in 27/28 dogs; the associated lymph node could not be found after amputation in one case (from a dog randomized to receive VSV). Two of the 27 cases evaluated had evidence of osteosarcoma metastasis in the lymph node, and the remaining 25 were negative for metastasis. One of the two dogs with lymph node metastasis at the time of amputation had shortened survival and was euthanized 64 days after surgery, with metastasis present throughout the abdomen and thorax on necropsy. Interestingly, the other dog with lymph node metastasis had prolonged survival and was still alive at the time of this report. Both cases with documented lymph node metastasis were treated with VSV.

### Virus pharmacokinetics

Monitoring viremia, virus shedding, and antiviral antibodies showed similar trends to previous studies evaluating intravenous VSV therapy in dogs.^14^ Viral RNA was detectable in whole blood samples at high quantities for the first 24 hours following infusion, with virus localizing primarily to PBMCs (Figure 5). Decay of viral RNA in blood coincided with an increase in detection of anti-VSV neutralizing antibodies, indicative of antibody-mediated virus clearance (Supplemental Figure S5). Infectious virus was detectable in PBMC samples at low quantities (below the limit of quantification) only at 1 hour post systemic VSV administration and no infectious virus was detected in PBMC samples collected after the 1h timepoint (data not shown). Virus shedding studies showed no detectable infectious virus in urine, rectal swab, or buccal swab samples, though viral RNA was detectable at low levels in some samples as shown (Supplemental Figure S6). Viremia, virus shedding, and neutralizing antibodies were measured in samples from two placebo-treated dogs showing no detectable viral RNA in blood or shedding samples, and no antiviral antibodies providing relevant negative controls in our pharmacokinetic analyses (data not shown). RNA from tumor specimens collected 10 days following VSV or placebo treatment showed detection of VSV RNA in bone tumor specimens from 2 of 22 VSV-treated dogs (supplemental table S4). The dose dependent efficacy of systemic VSV therapy^11^, the low amount and brief period that infectious virus was detectable, and the excellent safety of VSV infusion in this fixed-dose study collectively suggest higher doses of VSV-IFNβ-NIS can be safely administered in this treatment setting to potentially improve clinical response.

**Figure 5.**
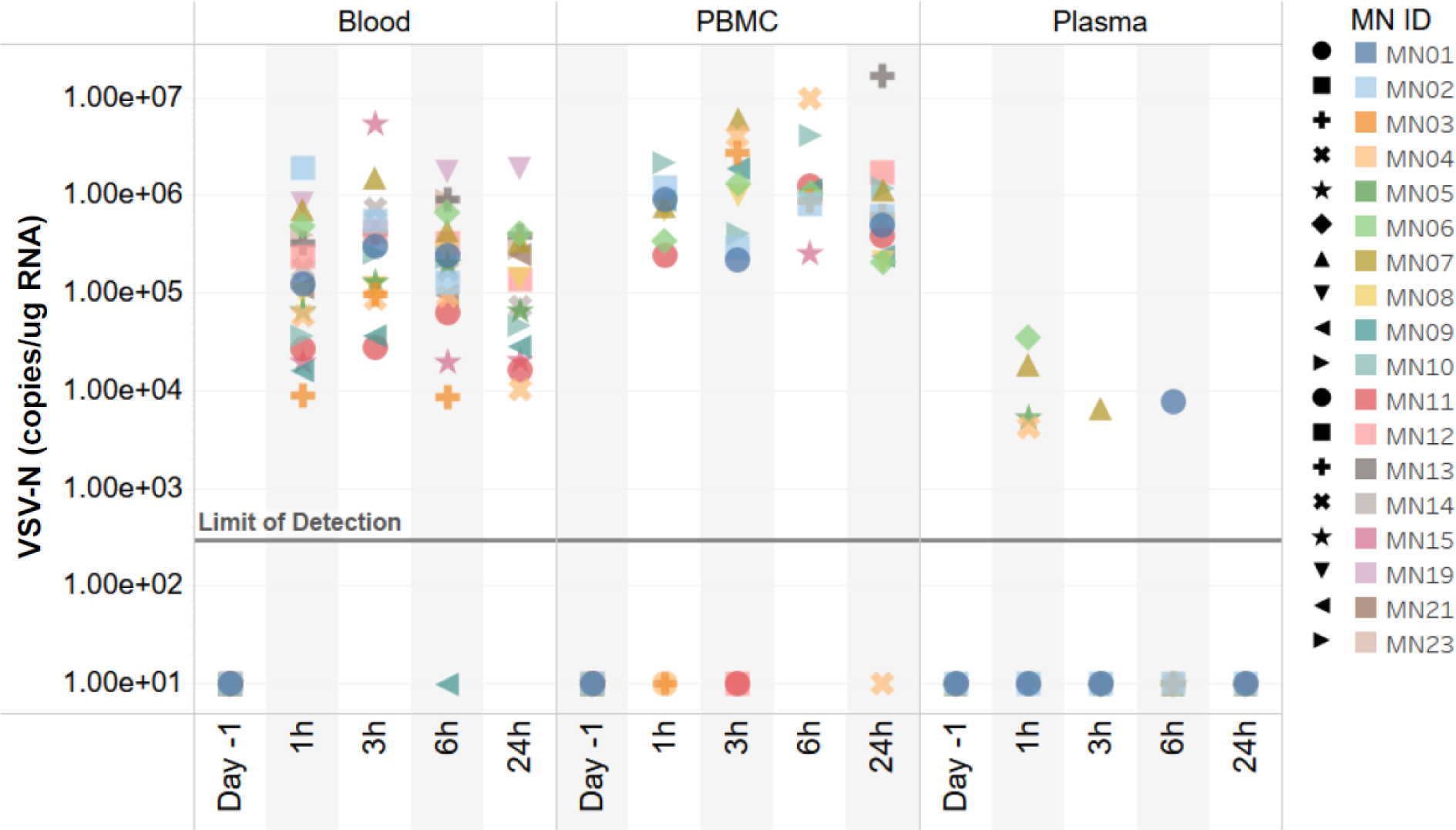
Acute viremia following systemic VSV infusion. Detection of VSV-N RNA copies in whole blood, PBMCs, and plasma samples collected at indicated time points following VSV infusion shows detection of viral RNA in blood and viral localization primarily to PBMCs.

### Pre-treatment immune infiltration is associated with longer overall survival

Acute cytokine responses were measured in the first 15 dogs enrolled and showed a transient elevation in several pro-inflammatory cytokines between 3 and 6 hours following VSV administration (Supplemental Figure S7). Three dogs that were long-term responders to VSV treatment, including the one diagnosed with hemangiosarcoma of bone, had high GM-CSF, IL-2, IL-7, and IL-15 levels at baseline.

Samples were examined for evidence of an immune component in the tumor response using next-generation RNA sequencing. Specifically, we used the gene cluster expression summary score (GCESS) method to summarize co-regulated gene clusters in tumor samples and their association with outcome.^20^ Pre- and post-treatment samples from primary osteosarcoma and metastatic osteosarcoma tumors grouped together, separate from both the cell lines derived from the pre-treatment tumors as well as from normal skin biopsy samples obtained from the same dogs at the time of amputation. Fourteen gene clusters were identified and were apparent in unsupervised hierarchical clustering heatmaps (Figure 6A-B), including a skin specific cluster (Figure 6C), as well as immune 1 (Figure 6D), immune 2 (Figure 6E), and cell cycle (Figure 6F) gene clusters previously identified in canine and human osteosarcoma.^20^ The distributions of cell cycle and immune GCESS across tumor samples, cell lines, and skin biopsy samples were as predicted, with no immune transcripts identified in osteosarcoma cell lines. VSV treatment did not notably impact the GCESS score of pre-versus post-treatment tumor specimens (Figure 6G-J). When we analyzed the association between GCESS and survival, we observed that the dogs with the highest pre-treatment immune 2 (T-cell) GCESS had longer survival times (Figure 7). This relationship was not observed for the CD37-positive monocyte-related GCESS, and somewhat surprisingly, it was also not evident for the cell cycle GCESS (Figure 7B-D). Upon comparison of the VIGOR cohort to a contemporary CCOGC cohort with available canine osteosarcoma RNAseq data (n=23, Supplemental Figure S8)^21^, we noted that a higher immune GCESS was associated with a longer progression-free survival in both groups, but that this effect was larger in the VSV-treated group. In the CCOGC cohort, all the dogs with longer survival had a higher T-cell anchored GCESS, but not all the dogs with higher CD8 GCESS had longer survival (Supplemental Figure S8C – see overlapping box plots); in the VIGOR group, all of the dogs with higher T-cell GCESS had longer survival (Figure 7E – see non-overlapping VSV high and VSV low box plots). The Kaplan-Meier Survival curve, showing increased overall survival in the VSV-treated dogs in the VIGOR group when compared with CCOGC cohort where RNA sequencing was available, is shown in Supplemental Figure S9.

**Figure 6.**
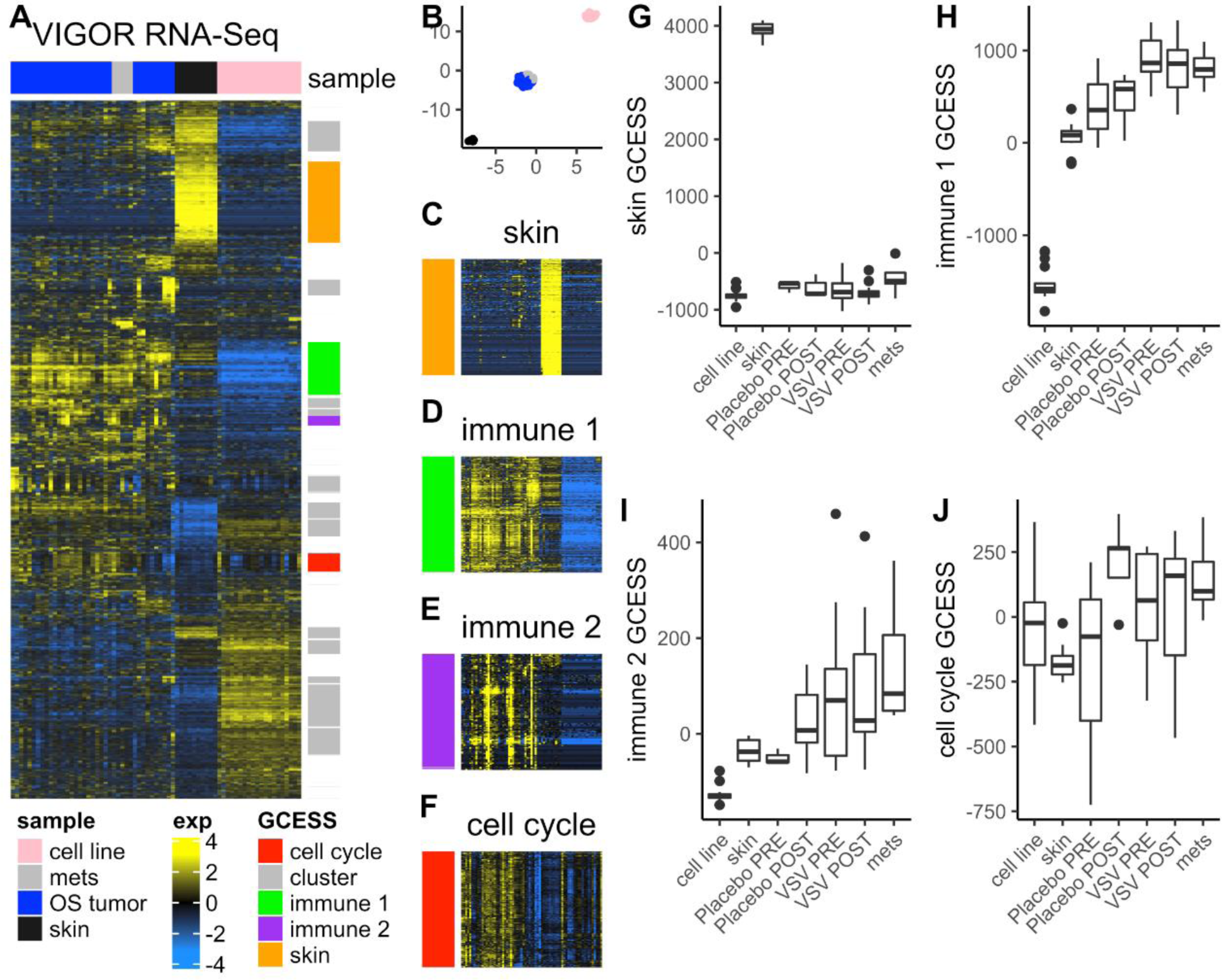
RNAseq analysis of canine tissues does not show significant changes immune gene signatures. (A) VIGOR RNAseq analysis of osteosarcoma tumor samples (pre- and post-treatment), lung metastases (where available from necropsy samples), skin biopsy, and osteosarcoma cell lines (derived from osteosarcoma tumors) clustered by tissue type. Data is log base 2 transformed mean centered and filtered for genes with high Standard deviation. Unsupervised hierarchical clustering resulted in 14 gene clusters including cell cycle, immune clusters and cluster composed primarily of genes expressed in skin samples. (B) UMAP projection of samples present in VIGOR dataset. Clear separation of OS tumors, skin and cell lines is apparently consistent with heatmap representation of the data. Samples are labeled with colors present in Figure 6A legend. (C-F) Zoomed in regions showing the C) skin specific cluster, D) immune 1 cluster composed of genes enriched in macrophage lineage immune genes, E) immune 2 cluster composed of genes enriched in T cell lineage immune genes and G) cell cycle enriched genes. (H-J) GCESS values generated by summing the genes present in each cluster for each sample are plotted in box plots representing samples grouped by sample type, treatment, and timepoint for H) skin specific GCESS, I) immune 1 GCESS, J) immune 2 GCESS, and F) Cell cycle GCESS. Unsupervised hierarchical clustering resulted in 14 gene clusters including cell cycle and immune clusters. GCESS scores were not significantly different in pre-vs. post-treatment VSV- and placebo treated tumor samples.

**Figure 7.**
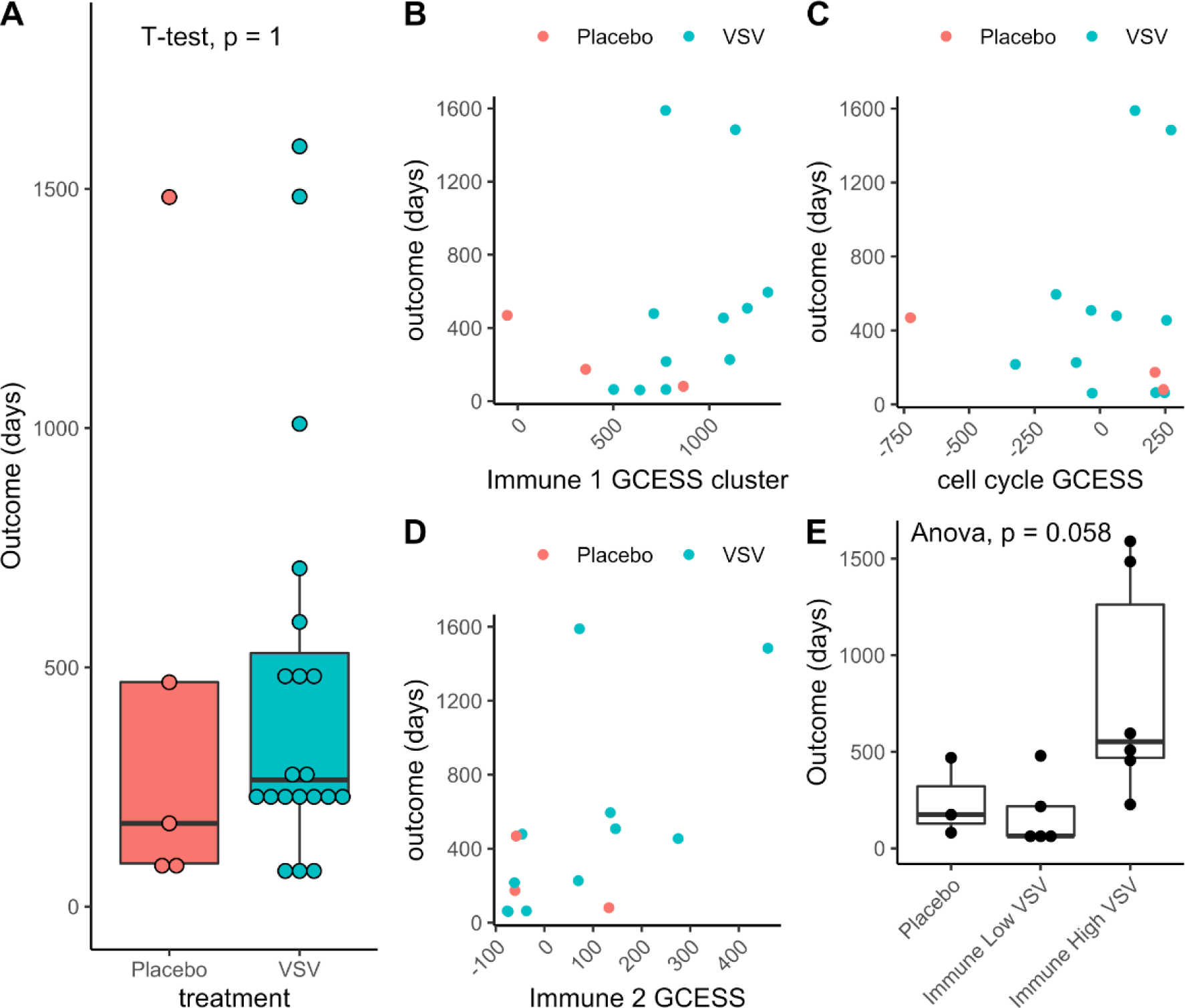
Survival outcomes correlate with pre-treatment T-cell GCESS. (A) Survival outcomes in placebo versus VSV treated dogs; (B-D) Plot of survival outcomes with (B) cell cycle GCESS, (C) immune 1 GCESS (macrophage/monocyte) (D), immune 2 GCESS (T-cell), indicates a correlation between survival and T-cell GCESS in pre-treatment tumor biopsy samples (E). The two dogs with non-osteosarcoma tumors of bone (hemangiosarcoma and rhabdomyosarcoma) and the dog that died of post-operative complications after the amputation surgery were excluded in this analysis.

## Discussion

Oncolytic viruses, including VSV, can selectively infect, replicate within, and kill tumor cells, promoting release of tumor associated antigens to drive robust anti-tumor immune responses.^7,22^ This approach is of particular interest in osteosarcoma, a genomically chaotic cancer, albeit with a low overall mutational burden and where recurrent mutations across the patient population are uncommon.^23–25^ Clinical studies evaluating oncolytic viruses to date have limited evaluation of the immune responses following treatment and highlight the need to identify and investigate biomarkers of clinical response.^26^ Here we describe the safety and efficacy of intravenous VSV-IFNβ-NIS therapy in naturally occurring canine appendicular osteosarcoma, a model utilized to inform development of new treatments for human osteosarcoma.^5^ Studies in spontaneous canine cancer supported clinical translation of VSV-IFNβ-NIS, a clinical-stage oncolytic virus that is being evaluated for the treatment of both solid tumors and hematologic malignancies.^8,9^ The major goals of this study were to establish the safety and clinical utility of VSV-IFNβ-NIS administered in the neoadjuvant setting in dogs with osteosarcoma, and to utilize correlatives studies in this spontaneous model to identify potential biomarkers of efficacy to oncolytic VSV therapy. Previous studies indicate that both safety and therapeutic response following systemic VSV therapy is dose dependent.^11,27^ In this study, we documented the safety profile of systemic therapy with VSV-cIFNβ-NIS in canine osteosarcoma. Intravenous VSV-cIFNβ-NIS, administered at 1×10^9^ TCID_50_/kg, was remarkably well-tolerated, with no clinically significant changes in any of the monitored parameters. Systemic viral distribution was documented with live virus detected (below the limit of quantification) in PBMCs within one-hour post-infusion, but not at later timepoints. Evidence of a systemic response was noted, as a subset of dogs examined showed acute elevations of inflammatory cytokines. No clinically significant adverse events were documented that were attributable to VSV therapy. The absence of adverse events paired with the low quantities and short duration of infectious virus detected in blood, suggest that the dose of systemic VSV therapy can be safely increased to potentially improve clinical outcomes.

We evaluated changes in GCESS and tumor inflammation scores on histopathology before and after treatment with VSV-IFNβ-NIS in this study. An increased tumor inflammation score (based on pathology assessment) was noted in tumor samples collected 10 days post VSV treatment compared to baseline samples. Clinical outcomes with neoadjuvant VSV therapy added to standard treatments for dogs with osteosarcoma were compared with two published control cohorts with dogs treated with only standard of care, including the NCI-COTC osteosarcoma cohort, a prospectively enrolled cohort designed to enable assessment and comparison of new osteosarcoma therapies in this clinically relevant model.^19^ While we did not observe a significant difference in survival between the groups, there was a tail of long-term survivors, defined as 75^th^ percentile overall survival from the NCI-COTC osteosarcoma cohort (479 days), noted in each population. In the VIGOR cohort, 35% of the dogs were considered long-term survivors compared to 25% and 26% in the NCI-COTC and UMN VMC control cohorts respectively. When we evaluated the association between immune GCESS and survival in the VIGOR cohort and CCOGC comparison group that had available RNAseq data, a higher T-cell anchored immune GCESS was associated with prolonged survival in both groups. However, this effect was more pronounced in the VSV-treated dogs. Our interpretation of these data is that the presence of a T-cell signature in osteosarcoma tumors prior to treatment increases the probability of longer survival, and the addition of VSV in the neoadjuvant setting enhances this effect.

In conclusion, these results show that systemically administered VSV-IFNβ-NIS can be used safely in the neoadjuvant setting to treat canine osteosarcoma. Systemic VSV-IFNβ-NIS is biologically active and can target the tumor microenvironment with evidence of increased tumor inflammation in VSV-treated osteosarcoma specimens. The intrinsic tumor immune status at baseline appears to influence the activity and therapeutic benefit of VSV, as dogs with higher immune 2 or T-cell anchored GCESS at baseline had prolonged survivals. The clinical safety and PK data, when viewed in the context of previous findings indicating dose dependent efficacy of systemic VSV therapy,^20^ suggest that there is scope to increase the systemic dose to improve clinical outcomes. Additionally, the observed therapeutic benefit in dogs with a pro-immune environment suggests that combinations with agents that promote immune infiltration or that amplify the immune response, such as immune checkpoint inhibitors, may further enhance the therapeutic benefits of VSV-IFNβ-NIS. The observed exceptional safety and immunological activity of VSV-IFNβ-NIS support further evaluation of neoadjuvant systemic VSV therapy including dose modification and combination therapy strategies for the treatment of sarcomas and translation to human cancer patients.

## Materials & Methods

### Manufacturing of VSV

Vesicular stomatitis virus (VSV) expressing canine interferon-beta (IFNβ) and the sodium iodide symporter (NIS), VSV-cIFNβ-NIS, was constructed as previously described.^10^ The VSV-cIFNβ-NIS lot used in veterinary studies was provided by the Mayo Clinic Viral Vector Production laboratory (VVPL). Virus product is stored at ≤ −65°C. Virus titers were determined by 50% tissue culture infective dose (TCID_50_) assays on Vero cells. The virus product was tested to confirm sterility (Mayo Clinic Department of Laboratory Medicine and Pathology) and absence of endotoxin using the LAL-Kinetic QCL Kit (Lonza). Whole genome sequencing was performed by utilizing reverse transcription of purified viral RNA to cDNA, polymerase chain reaction (PCR) of overlapping regions of viral genome, sequencing of PCR fragments (Eurofins Genomics), and assembly of sequencing results.

### *In vitro* efficacy of oncolytic VSV in canine osteosarcoma

Recombinant VSV vectors, including VSV expressing green fluorescent protein (GFP) or VSV expressing canine or human IFNβ and NIS (hIFNβ-NIS or cIFNβ-NIS), were used to infect three canine osteosarcoma cell lines, OSCA-78, OSCA-08, and OSCA-40, to evaluate *in vitro* efficacy of oncolytic VSV in canine osteosarcoma. Cells were grown in DMEM media containing 5% glucose and L-glutamine supplemented with 10% fetal bovine serum (FBS), 10mM HEPES buffer, and 0.1% Primocin, and cultured at 37°C in a humidified atmosphere of 5% CO_2_. Cell lines were authenticated based on short tandem repeats at regular intervals (IDEXX BioAnalytics). Virus titers were determined by TCID_50_ assay on BHK-21 cells at several timepoints after infection (12h, 24h, 36h, 48h, 60h, and 72h). Cell viability, as a percent of control, was determined by flow cytometry using a live/dead marker (Invitrogen) at several timepoints after infection (12h, 24h, 36h, 48h, 72h, and 96h). Cell lines are available through Kerafast, Inc (Boston, MA).

### Enrollment and eligibility criteria

Dogs with a diagnosis of appendicular bone sarcoma, based on radiographic appearance of the lesion and results of bone aspirate or biopsy, were considered for enrollment in the study (see CONSORT diagram, Supplementary Figure S1). All cases were enrolled at the University of Minnesota (UMN) College of Veterinary Medicine between June 2016 and January 2019. Screening diagnostics were performed prior to inclusion in the study, including physical examination, radiographs of the affected limb and thorax, general blood work (complete blood count, serum chemistry profile), computed tomography of the thorax and abdomen, and biopsy of the bone lesion. Inclusion criteria included cytologic or histologic diagnosis of sarcoma in an appendicular bone and body weight greater than 20kg (to facilitate safe collection of blood for the study). Exclusion criteria included evidence of metastatic lesions, evidence of pathologic fracture, intact reproductive status, clinically significant co-morbidities, treatment with other medications, and residence on a farm or other contact with farm animals (to reduce risk of VSV transmission to susceptible hosts).

### VSV dosing

Previous studies have demonstrated the safety of intravenous VSV-hIFNβ-NIS and VSV-cIFNβ-NIS therapy in dogs with cancer at doses up to 10^10^ TCID_50_/0.5m^2^, which converts to approximately 1×10^9^/kg.^14,27^ The VIGOR study enrolled a total of 28 dogs over 30 months (June 2016 to January 2019). The first 15 dogs received a single dose (1×10^9^ TCID_50_/kg) of neoadjuvant VSV-cIFNβ-NIS under an open label design. Because VSV at this dose was not associated with clinically significant adverse events and a preliminary survival benefit was noted, the next 13 dogs were randomized in a double-blinded fashion to receive a single dose of either intravenous VSV-cIFNβ-NIS or placebo (phosphate buffered saline solution (PBS)). The virus was diluted in sterile PBS to a volume of 10.5ml, which was administered by intravenous infusion over a period of 2-5 minutes. Dogs were housed for up to 24 hours in a BSL-2 housing facility. All procedures were carried out with approval of the University of Minnesota Institutional Animal Care and Use Committee (IACUC #1504-32486A and #1803-35759A) and Institutional Biosafety Committee (IBC #1504-32487H and # 1805-35967H).

### Standard of care therapy

All enrolled dogs were treated with standard of care therapy after treatment with intravenous VSV-cIFNβ-NIS or placebo. This consisted of amputation of the affected limb and removal of the associated lymph node(s), which occurred at day 10 following treatment, and initiation of chemotherapy approximately 14 days after amputation. Chemotherapy consisted of six cycles of carboplatin at a dose of 300 mg/m^2^, administered intravenously every 3 weeks, with standard monitoring for adverse chemotherapy effects.

### Biopsy procedures

Pre-treatment biopsies were obtained from the lesion using 4mm Michele trephine bone biopsy instruments. One pre-treatment tumor biopsy section was placed in 10% neutral buffered formalin for histopathology. A second pre-treatment section was further divided, with one piece used to establish a cancer cell line and a second piece frozen at −80°C for later RNA extraction. Ten days post-treatment with VSV-cIFNβ-NIS, dogs underwent surgical amputation of the affected limb and removal of the associated lymph node(s). A skin punch biopsy was performed, and the tissue frozen at −80°C for later RNA extraction. After amputation, the soft tissues of the limb were removed, and imaging of the affected bone was performed using a Faxitron (Supplementary Figure S2). The amputated limbs were sectioned longitudinally and tumor sections of approximately 5mm thickness were obtained and placed in 10% neutral buffered formalin. Tumor sections were placed in 10% ethanol for slow decalcification, or in acid for rapid decalcification. Separate sections of the tumor were frozen at −80°C for later RNA extraction. Cell lines derived as part of this study can be made available to the scientific community for research purposes under materials transfer agreements negotiated by the Regents of the University of Minnesota and Mayo Clinic.

### Monitoring virus pharmacokinetics

Assays to monitor viremia, virus shedding, and antiviral antibodies following intravenous VSV infusion were performed by Imanis Life Sciences using previously described protocols.^13,14,27^ Briefly, viremia was monitored by detection of infectious virus and viral RNA in blood samples collected at baseline and at specific time points after VSV treatment. For detection of viral RNA, whole blood was collected in RNAprotect animal blood tubes (Qiagen). For detection of infectious virus, peripheral blood mononuclear cells (PBMCs) were isolated from whole blood. Urine samples, fecal samples, and buccal swabs were collected after treatment with VSV-cIFNβ-NIS to test for presence of live infectious virus and viral RNA. Detection of infectious virus was performed by overlay of sample supernatants on susceptible BHK-21 cells. Viral RNA was detected in shedding and blood samples by quantitative reverse transcription-polymerase chain reaction (qRT-PCR) as previously described. Tumor specimens were stored at < −65°C at the time of surgical tumor resection (10 days post VSV therapy) for subsequent processing to isolate RNA and detect VSV RNA by qRT-PCR. Serum samples were obtained at baseline, and 3, 7, 14, 21, and 28 days post VSV infusion for detection of anti-VSV neutralizing antibodies.

### Serum cytokine monitoring

Serum samples were collected at baseline, 1, 3, 6 hours and 1-, 3-, 7- and 14-days post VSV infusion. Acute cytokine responses were determined as previously described by the COTC Pharmacodynamic Core at Colorado State University (CSU).^22^ The acute cytokines evaluated were: GM-CSF, IL-6, MCP-1, KC, IL-2, IP-10, IL-8, TNF-alpha, IL-10, IL-7, IL-1, and IL-18, all of the canine species.

### Clinical monitoring

Body temperature was monitored after treatment with VSV-cIFNβ-NIS, every 3 hours for the first 24 hours following treatment, then daily. Blood work to evaluate alanine aminotransferase (ALT), aspartate aminotransferase (AST), total white blood cell count, lymphocyte count, and clotting times (PT and aPTT) was completed at baseline, and at days 3, 7, 14, 21, and 28 after treatment.

### Histopathology

Pre-treatment core tumor biopsies and post-treatment amputation-resected tumor samples were evaluated with H&E staining to obtain a definitive histologic diagnosis. Samples were also scored based on tumor inflammation, fibrosis, and necrosis. Inflammation was graded on a 4-point scale, where 0 = absence of inflammation, 1 = minimal inflammation, 2 = mild inflammation, 3 = moderate inflammation, and 4 = marked inflammation. Fibrosis was graded on a 4-point scale, where 0 = absence of fibrosis, 1 = minimal fibrosis, 2 = mild fibrosis, 3 = moderate fibrosis, and 4 = marked fibrosis. Necrosis was graded on a 3-point scale, where 0 = no necrosis, 1 = < 15%, 2 = 15 - 50%, and 3 = > 50% of the section had apparent necrosis. The draining lymph node was evaluated for presence or absence of metastatic osteosarcoma cells. During the placebo-arm of the study, the pathologist was blinded to the treatment assignment of cases (VSV or placebo).

### Collection of PBMCs for immune monitoring

Blood samples were obtained for isolation of PBMCs and plasma for immune monitoring. Blood was collected in a commercially available tube for PBMC isolation (BD Mononuclear Cell Preparation Tube), and the manufacturer’s protocol was followed. PBMCs were frozen in FBS with 10% DMSO and stored in liquid nitrogen until use.

### Diagnostic imaging

Cases were monitored with diagnostic imaging to evaluate for metastatic disease throughout the study period. Thoracic radiographs were performed at study evaluation (d-1), and on days 60, 120, 180, and 360 post-treatment. Computed tomography (CT) was performed at study evaluation (d-1) and at day 90 post-treatment.

### RNA extraction from tumor specimens

VIGOR: RNA was extracted from tumor tissue at baseline, and from tumor tissue samples obtained at amputation, post-treatment. RNA was also extracted from cell lines that were established pre-treatment from each case, and from normal skin tissue from each case as a non-malignant control. When available, RNA was also extracted from lung metastases. Tissue samples were disrupted with a tissue homogenizer, and RNA was extracted using a commercially available kit (QIAGEN RNeasy mini kit). RNA was stored at −80°C until submission to the University of Minnesota Genomics Center (UMGC).

CCOGC: A comparison population (n = 23), consisted of tissue samples provided by the Pfizer CCOGC Biospecimen Repository. Samples with a definitive, histological diagnosis of osteosarcoma were selected for RNA sequencing based on the availability of sufficient frozen tissue (>50 mg), treatment information, RNA integrity numbers (RIN) >6, and available follow-up information. These were included in the final selection for next-generation RNA sequencing.

### Library preparation and next-generation sequencing

Unique dual-indexed sequencing libraries were prepared using the Clontech Stranded Total RNA-Seq Kit v2 - Pico Input Mammalian kit. RNA sequencing (2×50-bp paired-end, on a NovaSeq S2) was performed at the UMGC. Approximately 3,100M pass filter reads were generated for the run. Mean quality scores were above Q30 for all libraries. Successful libraries were generated from 64 samples (15 pre-treatment tumors, 20 post-treatment tumors, 18 cancer cell lines generated from pre-treatment tumors, and 11 skin samples) from 24 dogs. Additionally, two dogs had pulmonary metastatic lesions obtained at necropsy from which libraries were generated from tissue and from cell lines. A total of 23 osteosarcoma tissue samples were included in the CCOGC dataset.^21^

## Gene cluster expression summary score (GCESS) and Bioinformatics analysis

Initial quality control analysis of RNA sequencing FASTQ data was performed using FastQC software (v0.11.5). FASTQ data were trimmed with Trimmomatic (v0.33.0). Kallisto (v0.43.0) was used for pseudoalignment and quantifying transcript abundance. Sequencing reads were aligned to the canine reference genome (CanFam3.1). Transcript abundance counts were generated, and quantile normalized to correct for differences in sequence counts. Gene cluster expression summary score, or GCESS, analyses were performed as previously described on average linkage hierarchical clustered data.^20^ Briefly, clusters of genes are identified based on patterns of correlated expression, such as those associated with immune cell infiltration or cell cycling. After mean-centering and log_2_-transformation, individual gene values in each cluster are added together resulting in a single summary score for the cluster that is reflective of the overall degree of gene expression. UMAP was used to generate a 2-dimensional representation of the transcriptional patterns present in the dataset.^28^

### Outcome assessment

Formal clinical follow up, including physical examinations, blood work, and diagnostic imaging, occurred for one year after treatment with VSV-cIFNβ-NIS. Thereafter, informal follow up occurred until relapse or death. Necropsies were obtained, when possible, with owner consent. Outcomes assessed included event-free survival (EFS, time to relapse) and overall survival (time to death). The dogs enrolled in the VIGOR study (n = 28) were compared with three control populations. The first population consisted of dogs with appendicular osteosarcoma treated at the same institution (UMN, n = 57) between January 2006 and December 2018. The second population consisted of the control arm of an NCI Comparative Oncology Trials Consortium multicenter clinical trial in dogs with appendicular osteosarcoma (COTC, n = 157) enrolled between November 2015 and February 2018.^13^ Both comparison populations were treated with standard of care therapy, consisting of amputation of the affected limb and carboplatin chemotherapy administered every 21 days at 300mg/m^2^. A third comparison population (CCOGC, n = 23), consisted of dogs treated with standard of care (chemotherapy and amputation) with known outcome data, enrolled in a multicenter study between April 2008 and March 2011 for which samples were available for tumor next-generation RNA sequencing.^21^ This population was used for comparison of survival and tumor gene expression.

### Statistical analyses

Tumor inflammation scores were compared between pre-treatment and post-treatment tumor samples using paired student’s t-test. P-values are reported without inference to significance, as recommended by the American Statistical Association.^23^ Kaplan-Meier survival curves were generated for the dogs treated with VSV-cIFNβ-NIS in the VIGOR study and the comparison populations. The EFS was calculated as the time between limb amputation and the first detection of metastasis. Dogs were censored if they did not have metastatic disease at the time of last follow-up. Overall survival was calculated as the time between limb amputation and death. Dogs were censored from survival analyses if they were found to have tumors of bone that were not osteosarcoma, if they died of non-osteosarcoma related causes, or if they were alive at last follow up.

## Author contributions

SN, SJR, KMS, and JFM were involved in overall study design and execution including obtaining regulatory approvals and funding to implement the described studies. AC, DG, KB, SP, AW, and KMM were responsible for sample collection and processing. AC, KMS, AB, and MSH were responsible for clinical follow-up. LS and SN oversaw and executed correlative pharmacokinetic studies (assays to measure viremia, virus shedding, antibodies, etc.). CP assisted in data processing and analysis. ALS and LJM analyzed sequencing data. MDS and MW completed methodological design for gross sample collection, imaging with the Faxitron, and gross lesion identification, assessment, and sampling for optimal downstream analysis. MW organized pathological images, maintained supplies, and facilitated communication regarding samples. IC, AFT, and MGOS were responsible for analyzing histopathology specimens. SN, ALS, LJM, JFM, and KMM were responsible for additional data analysis. GRC and JSK were responsible for statistical design and analysis. SN, KMM, ALS, and JFM were responsible for data compilation and manuscript preparation. All authors were responsible for manuscript review and revision.

## Data availability statement

RNA-Seq expression profiling data from canine osteosarcoma tumor samples obtained from Canine Comparative Oncology and Genomics Consortium (CCOGC) and VIGOR cohorts are submitted and available to the public via Gene Expression Omnibus (GEO), a public functional genomics data repository. CCOGC dataset GEO accession number: GSE239948. VIGOR dataset GEO accession number: GSE240033.

## Acknowledgements

The authors would like to acknowledge Milcah Scott for assistance in training, including cell culture and sequencing techniques. The authors would like to acknowledge Mitzi Lewellen for oversight of sample databases. The authors would also like to acknowledge Katalin Kovacs and Paula Overn for their assistance in preparation of histology specimens. The authors acknowledge the Clinical Investigation Center (CIC) of the University of Minnesota College of Veterinary Medicine, the Veterinary Diagnostic Laboratory (VDL) of the University of Minnesota, the University of Minnesota Genomics Center (UMGC), and the Minnesota Supercomputing Institute (MSI) at the University of Minnesota for providing resources that contributed to the research results reported within this paper. The authors wish to thank the VDL Pathologists for their assistance in sample preparation, including Dr. Arno Wuenschmann, Dr. Erik Olson, and Dr. Jaclyn Dykstra. The authors also wish to thank the CCOGC, Inc. for providing a set of control tumor samples for this study, as well as Dr. Subbaya Subramanian for help with procuring the CCOGC samples. Additionally, the authors wish to thank Dr. Amy LeBlanc and Christina Mazcko from the NCI COTC for providing the raw survival data for the control cohort treated with the standard of care.

The authors wish to thank the clinical teams at the University of Minnesota College of Veterinary Medicine Lewis Small Animal Teaching Hospital for the clinical care of enrolled dogs, including Oncology, Soft Tissue Surgery, Diagnostic Imaging, the Veterinary Diagnostic Laboratory, Clinical Pathology, and associated support staff.

## Funding

The work described in this manuscript was supported by grant MNP #15.25 from the Minnesota Partnership for Biotechnology and Medical Genomics, sponsored research from Vyriad, Inc. (Rochester MN), grant R21 CA208529 from the National Cancer Institute (NCI) of the National Institutes of Health (NIH), United States Public Health Service, and grants CA170218 and CA190276 from the US Department of Defense Peer Reviewed Cancer Research Program. NCI Comprehensive Cancer Center Support Grant P30 CA077598, to the Masonic Cancer Center, University of Minnesota, provided support for genomics and bioinformatics. Tumor samples and relevant metadata from the CCOGC cohort were provided *pro bono* by the CCOGC, Inc. KMM was supported in part by institutional training grant in Molecular, Genetic, and Cellular Targets of Cancer T32 CA009138 from the NCI. ALS was supported in part by grant R50 CA211249 from the NCI. JFM is supported in part by the Alvin and June Perlman Endowed Chair in Animal Oncology. The authors acknowledge support from individual donors to the Animal Cancer Care and Research Program, University of Minnesota. The content of this manuscript is solely the responsibility of the authors and does not necessarily represent the official views of any of the funding agencies listed above.

## Conflict of Interest

Drs S. Naik, S.J. Russell, and Mayo Clinic have a financial conflict of interested related to this research. S. Naik and SJ Russell are inventors on patents related to oncolytic Vesicular stomatitis virus that has been licensed by Mayo Clinic to Vyriad. S. Naik and S.J. Russell have equity interests in Vyriad, and S.J. Russell is the CEO of Vyriad. The research has been reviewed by the Mayo Clinic Conflict of Interest Review Board and is being conducted in compliance with Mayo Conflict of Interest policies.

## Notes

### Summary of Updates

Title was revised to more accurately reflect study findings. Abstract and Introduction were revised to increase emphasis on the importance of studies testing oncolytic viruses in the neoadjuvant setting. Changes were made to Figure 6 and 7 to improve layout. Results section updated to provide more detail on experimental results and reflect changes made to figures 6 and 7. Data availability statement and keywords were added.

