## Supplemental Data for "Neoadjuvant systemic oncolytic vesicular stomatitis virus is safe and may enhance long-term survivorship in dogs with naturally occurring osteosarcoma"

**Table S1. List of adverse events that occurred following VSV administration were uncommon and considered unlikely to be related to treatment.**

Dogs were monitored closely for AEs following systemic VSV administration. AEs were uncommon and those that occurred were considered unlikely to be related to VSV treatment.

| Dog ID | Adverse Events (AEs) | Interval (days) from VSV infusion to AE | AE grade | AE related to VSV? | AE related to disease? | AE outcome |
|---|---|---|---|---|---|---|
| MN01 | Facial nerve paralysis | 106 | 1 | Unlikely | Unlikely | Not recovered |
| MN03 | Pneuomonia/ intrapulmonary hemorrhage | 142 | 1 | Unlikely | Likely[1] | Recovered |
| MN17 | Hypovolemic shock after surgery | 5 | 5 | Unlikely | Possible | Death |

[1]Although aspiration pneumonia was the presumptive diagnosis, the sum of the data from this case suggests that the dog had intrapulmonary hemorrhage from one of its many pulmonary nodules, and not pneumonia. The hemorrhage resolved, but the nodules did not, and the dog eventually died from progressive disease. While the cause of hemorrhage cannot be clearly ascertained, it was most likely due to disease progression.

**Table S2. Clinical information and survival data for enrolled dogs.** Censored events due to death from other causes are noted in red font, and animals that are still alive at the time of analysis are noted in blue font.

| Case # | Breed | Weight (kg) | Age (years) | Tumor location | TX | Status at study end | Presumed cause of death | EFS (days) | Overall survival (days) |
|---|---|---|---|---|---|---|---|---|---|
| MN01 | Golden Retriever | 30 | 6 | L proximal humerus | VSV | Dead | Disease | 518 | 595 |
| MN02 | Rottweiler | 59 | 7 | L distal radius | VSV | Dead | Other cause | 356 | 707 |
| MN03 | Boxer | 25.7 | 7 | L distal radius | VSV | Dead | Disease | 133 | 227 |
| MN04 | Mixed Breed | 35 | 11 | L carpus | VSV | Dead | Other cause | 1009 | 1009 |
| MN05 | Australian Shepherd | 20.2 | 7 | R proximal humerus | VSV | Dead | Disease | 273 | 275 |
| MN06 | Mixed Breed | 50 | 5 | R distial tibia | VSV | Dead | Disease | 64 | 64 |
| MN07 | Mixed Breed | 42 | 2 | L proximal humerus | VSV | Dead | Disease | 180 | 243 |
| MN08 | Labrador Retriever | 52 | 9.5 | L proximal humerus | VSV | Dead | Disease | 355 | 508 |
| MN09 | Golden Retriever | 33.7 | 8 | L distal radius | VSV | Dead | Disease | 181 | 237 |
| MN10 | Mixed Breed | 31.2 | 3 | R distal radius * | VSV | **Alive** | | 1683 | 1683 |
| MN11 | Golden Retriever | 29.9 | 9.5 | R proximal humerus | VSV | Dead | Disease | 194 | 455 |
| MN12 | Old English Sheepdog | 34.7 | 9 | L distal femur | VSV | Dead | Disease | 82 | 205 |
| MN13 | Golden Retriever | 33.8 | 9.5 | R distal tibia | VSV | Dead | Disease | 169 | 277 |
| MN14 | German Shorthair Pointer | 30 | 8 | L distal ulna | VSV | Dead | Disease | 50 | 88 |
| MN15 | Greyhound | 29.7 | 8 | L proximal tibia | VSV | **Alive** | | 1589 | 1589 |
| MN16 | Labrador Retriever | 30.5 | 8.5 | L distal radius | Placebo | Dead | Disease | 60 | 81 |
| MN17 | Irish Wolfhound | 37.4 | 6 | L distal tibia | VSV | Dead | Surgical complications | 0 | 0 |
| MN18 | Mixed Breed | 26.3 | 9 | R femoral diaphysis | Placebo | **Alive** | | 1483 | 1483 |
| MN19 | Mixed Breed | 26.9 | 9 | R proximal humerus | VSV | **Alive** | | 1484 | 1484 |
| MN20 | Mastiff | 81.6 | 2 | R proximal humerus ** | Placebo | Dead | Disease | 53 | 74 |
| MN21 | St. Bernard | 50.6 | 7 | L radius | VSV | Dead | Disease | 49 | 61 |
| MN22 | Great Pyrenees | 39.3 | 5.5 | L distal radius | Placebo | Dead | Disease | 63 | 174 |
| MN23 | Labrador Retriever | 26.3 | 12.5 | R proximal humerus | VSV | Dead | Disease | 81 | 237 |
| MN24 | German Shepherd | 48.4 | 9 | R proximal humerus | Placebo | Dead | Disease | 56 | 90 |
| MN25 | Mixed Breed | 30.8 | 5 | L proximal tibia | VSV | Dead | Disease | 171 | 217 |
| MN26 | Mastiff | 73.4 | 8 | R proximal humerus | Placebo | Dead | Disease | 377 | 476 |
| MN27 | Vizsla | 24.8 | 8 | R distal tibia | VSV | Dead | Disease | 54 | 254 |
| MN28 | Golden Retriever | 23.6 | 12 | R distal radius | VSV | Dead | Other cause | 479 | 479 |

**Table S3. Pathology assessment of pre- and post-treatment tumor specimens**

FFPE tissues from pre-treatment (pre-tx) tumor biopsies and tumors resected 10 days after VSV or placebo administration (post-tx) were assessed by a veterinary pathologist who was blinded to treatment group. Tumors were score for inflammation, fibrosis, and necrosis. N/E indicates non-evaluable specimens.

| | | Intratumoral Inflammation (Grade) | | Intratumoral Fibrosis (Grade) | | Intratumoral Necrosis (Grade) | | | |
|---|---|---|---|---|---|---|---|---|---|
| **Case #** | **TX** | **Pre-tx** | **Post-tx** | **Pre-tx** | **Post-tx** | **Pre-tx** | **Post-tx** | **Micro-necrosis Present** | **Tumor Grade** |
| MN01 | VSV | 0 | 0 | 0 | 0 | n/e | 1 | No | 3 |
| MN02 | VSV | n/e | 3 | 0 | 4 | n/e | 4 | Yes | 3 |
| MN03 | VSV | 1 | 3 | 1 | 0 | 2 | 4 | No | 3 |
| MN04 | VSV | 0 | 2 | 0 | 3 | 0 | 4 | No | 3 |
| MN05 | VSV | n/e | 2 | 0 | 3 | n/e | 4 | Yes | 3 |
| MN06 | VSV | n/e | 0 | n/e | 3 | n/e | 4 | No | 3 |
| MN07 | VSV | n/e | 1 | n/e | 3 | n/e | 3 | Yes | 3 |
| MN08 | VSV | n/e | 0 | 2 | 4 | n/e | 4 | No | 2 |
| MN09 | VSV | n/e | 0 | n/e | 3 | n/e | 3 | No | 3 |
| MN10 | VSV | 0 | 3 | 2 | 3 | n/e | n/e | No | N/A |
| MN11 | VSV | 0 | 4 | 0 | 0 | 0 | 4 | No | 3 |
| MN12 | VSV | 0 | 2 | 2 | 3 | n/e | 4 | Yes | 3 |
| MN13 | VSV | n/e | 2 | n/e | 2 | n/e | 4 | No | 3 |
| MN14 | VSV | 1 | 2 | 3 | 3 | n/e | 4 | Yes | 3 |
| MN15 | VSV | n/e | 1 | 0 | 0 | n/e | 4 | Yes | 3 |
| MN16 | Placebo | 0 | 0 | 0 | 1 | n/e | 4 | Yes | 3 |
| MN17 | VSV | 0 | 1 | 0 | 3 | 0 | 0 | Yes | 1 |
| MN18 | Placebo | 0 | 2 | 0 | 0 | n/e | 3 | Yes | 3 |
| MN19 | VSV | 0 | 3 | 0 | 3 | n/e | 3 | Yes | 3 |
| MN20 | Placebo | 0 | 0 | 0 | 2 | n/e | 4 | No | 3 |
| MN21 | VSV | n/e | 3 | 0 | 3 | n/e | 2 | Yes | 2 |
| MN22 | Placebo | 0 | 2 | 1 | 0 | 3 | 4 | No | 3 |
| MN23 | VSV | n/e | 0 | 0 | 2 | n/e | 3 | Yes | 3 |
| MN24 | Placebo | n/e | 0 | 0 | 2 | n/e | 3 | No | 3 |
| MN25 | VSV | 0 | 0 | 1 | 0 | n/e | 4 | No | 3 |
| MN26 | Placebo | n/e | 0 | 0 | 0 | n/e | 4 | No | 2 |
| MN27 | VSV | 0 | 0 | 0 | 2 | n/e | 4 | No | 3 |
| MN28 | VSV | 0 | 0 | 2 | 2 | 0 | 2 | No | 2 |

**Table S4. Detection of VSV RNA in tumor specimens.**

Detection of VSV-N RNA copies in RNA isolated from resected bone tumor specimens collected 10 days following VSV or placebo administration. Lung metastasis specimens during a necropsy was performed on one dog (MN25) that was humanely euthanized due to disease progression and similarly analyzed for detection of VSV RNA.

| Dog ID | Treatment group | Tumor specimen | VSV-N (copies/ug RNA) |
|---|---|---|---|
| MN01 | VSV | Bone tissue RNA (Day 10 specimen) | BLD |
| MN02 | VSV | Bone tissue RNA (Day 10 specimen) | BLD |
| MN03 | VSV | Bone tissue RNA (Day 10 specimen) | BLD |
| MN04 | VSV | Bone tissue RNA (Day 10 specimen) | BLD |
| MN05 | VSV | Bone tissue RNA (Day 10 specimen) | BLD |
| MN06 | VSV | Bone tissue RNA (Day 10 specimen) | BLD |
| MN07 | VSV | Bone tissue RNA (Day 10 specimen) | BLD |
| MN08 | VSV | Bone tissue RNA (Day 10 specimen) | BLD |
| MN09 | VSV | Bone tissue RNA (Day 10 specimen) | BLD |
| MN10 | VSV | Bone tissue RNA (Day 10 specimen) | BLD |
| MN11 | VSV | Bone tissue RNA (Day 10 specimen) | BLD |
| MN12 | VSV | Bone tissue RNA (Day 10 specimen) | BLD |
| MN13 | VSV | Bone tissue RNA (Day 10 specimen) | BLD |
| **MN14** | **VSV** | **Bone tissue RNA (Day 10 specimen)** | **2.74E+03** |
| MN15 | VSV | Bone tissue RNA (Day 10 specimen) | BLD |
| MN17 | VSV | Bone tissue RNA (Day 10 specimen) | BLD |
| MN19 | VSV | Bone tissue RNA (Day 10 specimen) | BLD |
| MN21 | VSV | Bone tissue RNA (Day 10 specimen) | BLD |
| MN23 | VSV | Bone tissue RNA (Day 10 specimen) | BLD |
| **MN25** | **VSV** | **Bone tissue RNA (Day 10 specimen)** | **2.62E+05** |
| MN25 | VSV | Lung metastasis #1 (necropsy specimen) | BLD |
| MN25 | VSV | Lung metastasis #2 (necropsy specimen) | BLD |
| MN27 | VSV | Bone tissue RNA (Day 10 specimen) | BLD |
| MN28 | VSV | Bone tissue RNA (Day 10 specimen) | BLD |
| MN16 | Placebo | Bone tissue RNA (Day 10 specimen) | BLD |
| MN18 | Placebo | Bone tissue RNA (Day 10 specimen) | BLD |
| MN20 | Placebo | Bone tissue RNA (Day 10 specimen) | BLD |
| MN22 | Placebo | Bone tissue RNA (Day 10 specimen) | BLD |
| MN24 | Placebo | Bone tissue RNA (Day 10 specimen) | NA |
| MN26 | Placebo | Bone tissue RNA (Day 10 specimen) | BLD |

**Figure S1.** VIGOR study CONSORT diagram showing screening, enrollment, treatment, and follow-up assessments.

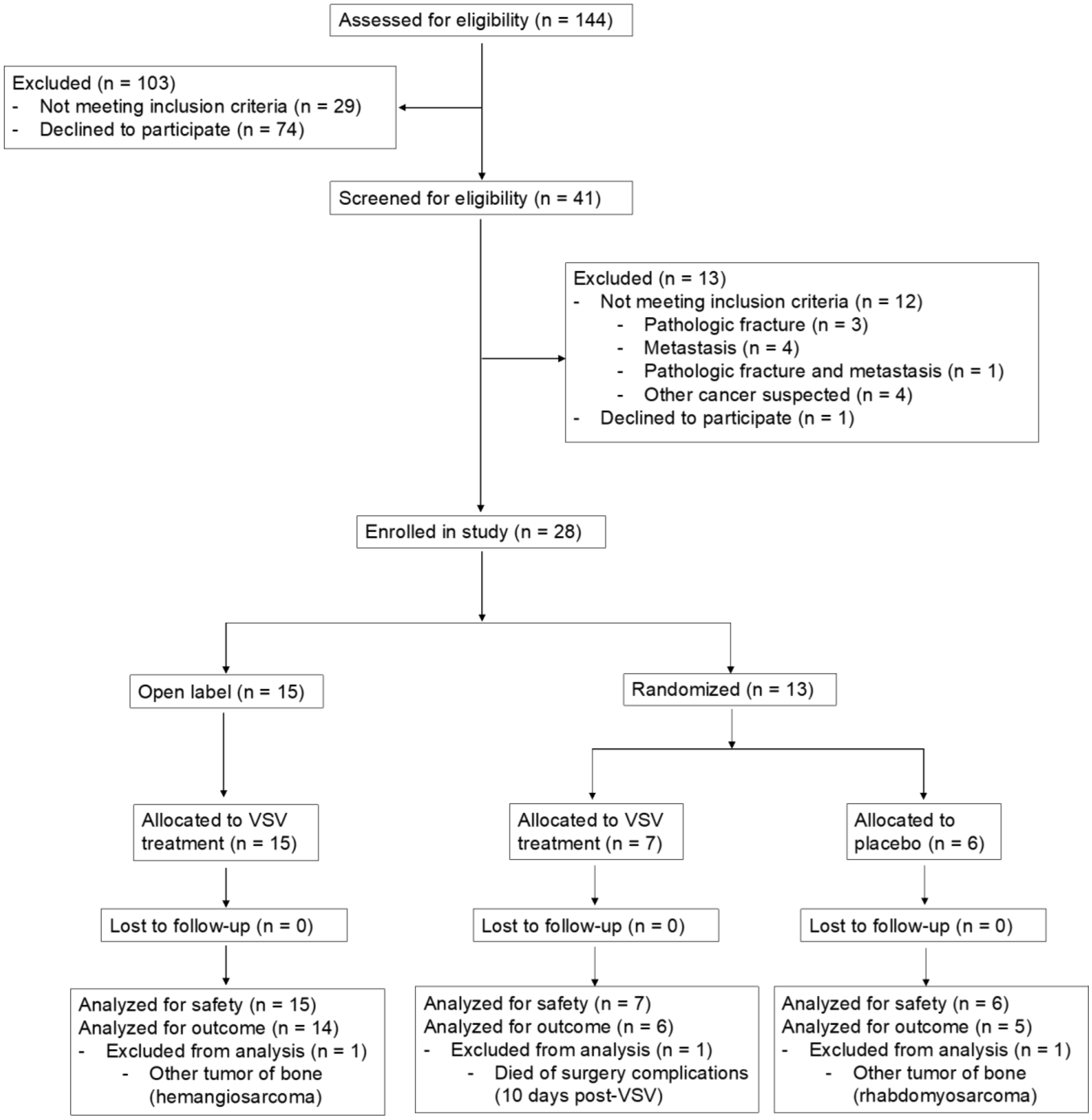

**Figure S2. Faxitron imaging after amputation allows for more accurate tumor size determination and guides tumor sample collection.** (A) Radiographic image of a left proximal humeral osteosarcoma lesion, and (B) Faxitron image and gross pathology evaluation of the same lesion. White dashed lines indicate tumor borders on visual inspection.

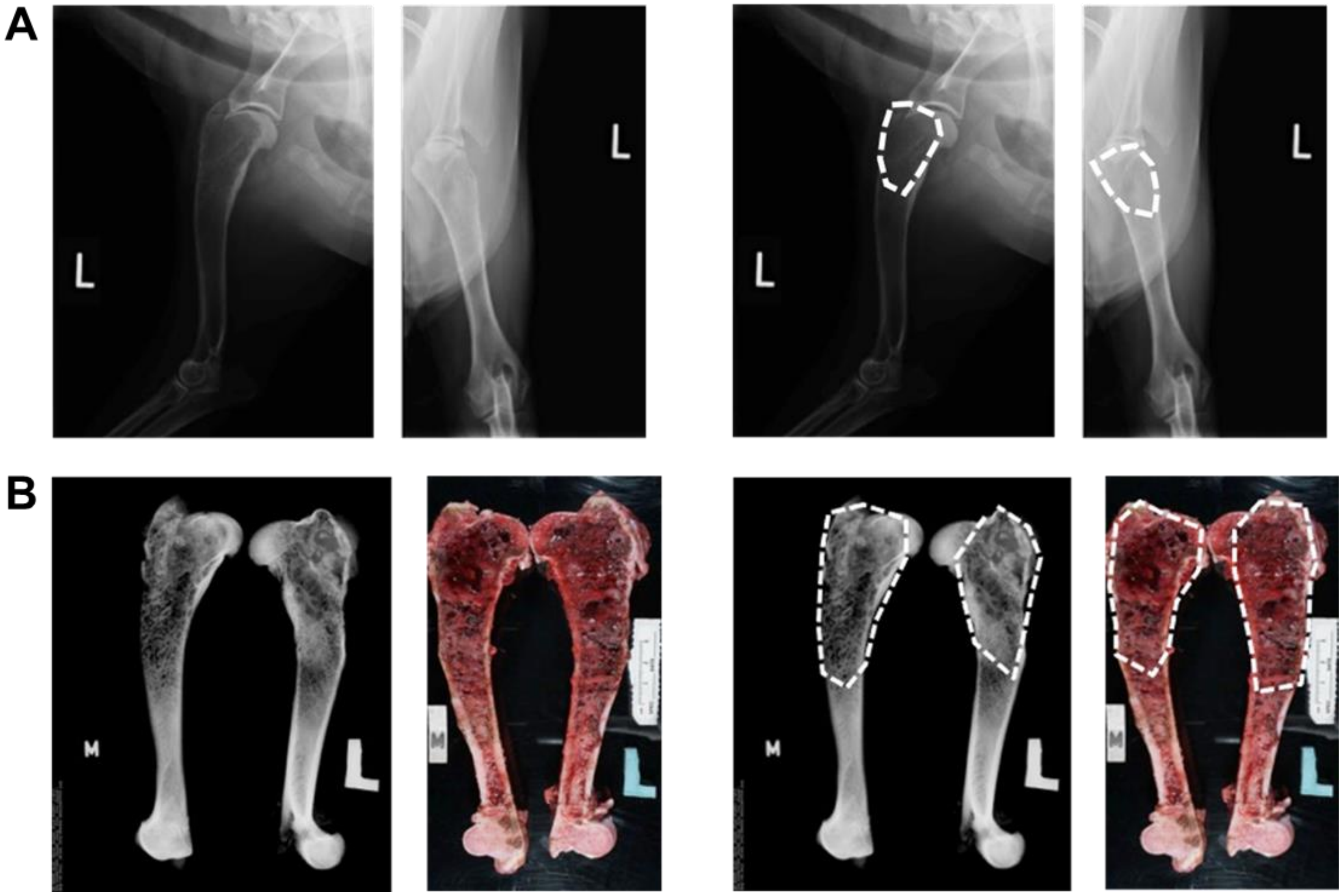

**Figure S3. Administration of systemic vesicular stomatitis virus (VSV) in dogs with osteosarcoma did not cause any clinically significant laboratory abnormalities.** Common laboratory tests including blood chemistry and complete blood count (CBC) were used to monitor clinical safety following VSV infusion showing no significant abnormalities relative to baseline. Blood test results from the first 11 enrolled dogs are shown indicating no significant changes in (A) liver function testes including alanine aminotransferase (ALT) and aspartate aminotransferase (AST); and (B) lymphocytes (LYMPH) and total white blood cell count (WBC).

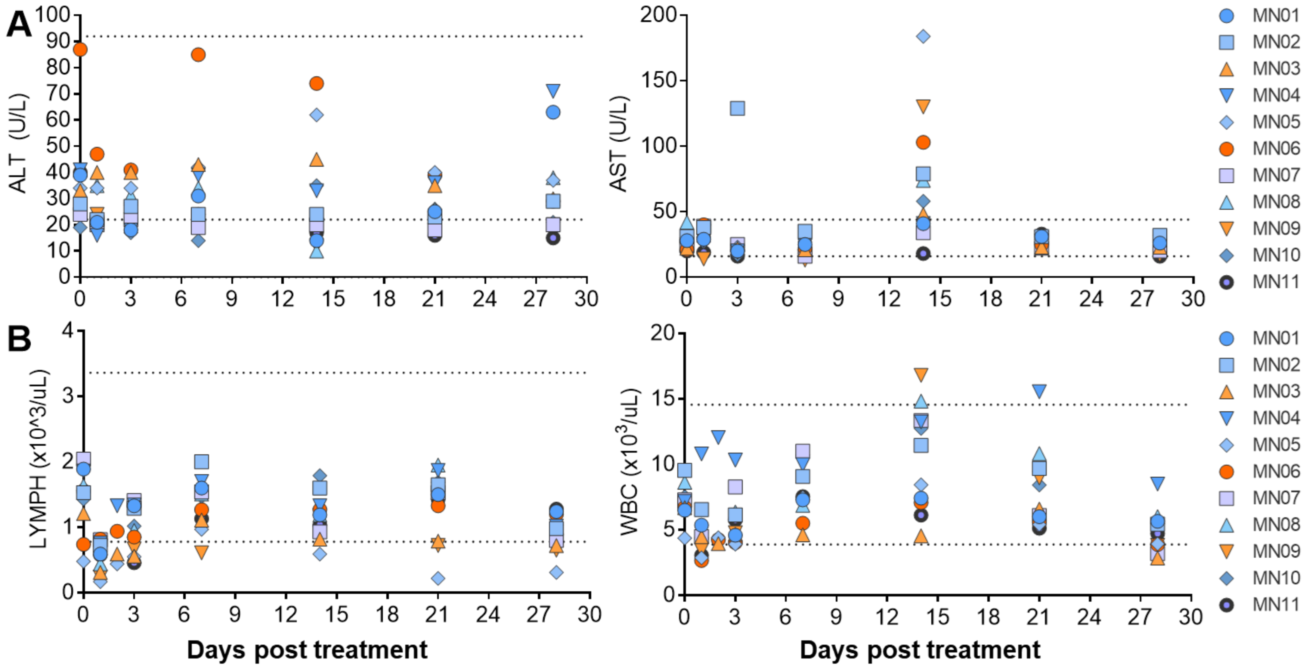

**Figure S4. Small foci of micronecrosis seen in a subset of cases treated with systemic VSV may indicate areas of viral oncolysis.** H&E histopathology of amputation samples obtained 10 days after treatment with systemic VSV. (A) Large areas of necrosis commonly associated with ischemia observed in resected osteosarcoma specimens. (B) Small focal areas of micronecrosis were seen in a subset of treated cases, potentially due to viral infection of tumor cells and subsequent cell lysis.

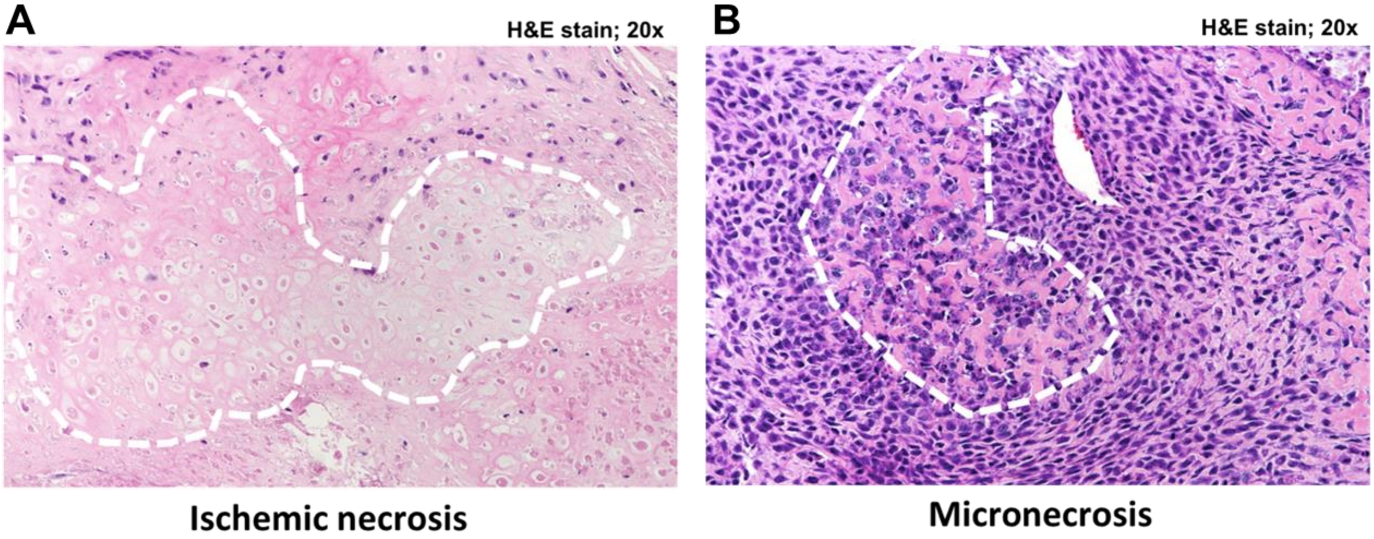

**Figure S5. Virus clearance from blood corresponds with generation of anti-VSV neutralizing antibodies**

(A) qRT-PCR detection of VSV-N copies in PBMC samples and (B) Neutralizing antibody titers in serum samples collected from a subset of dogs that received systemic VSV therapy. All evaluated dogs developed neutralizing antibodies at or above the limit of detection (LOD) within approximately 7 days after VSV administration.

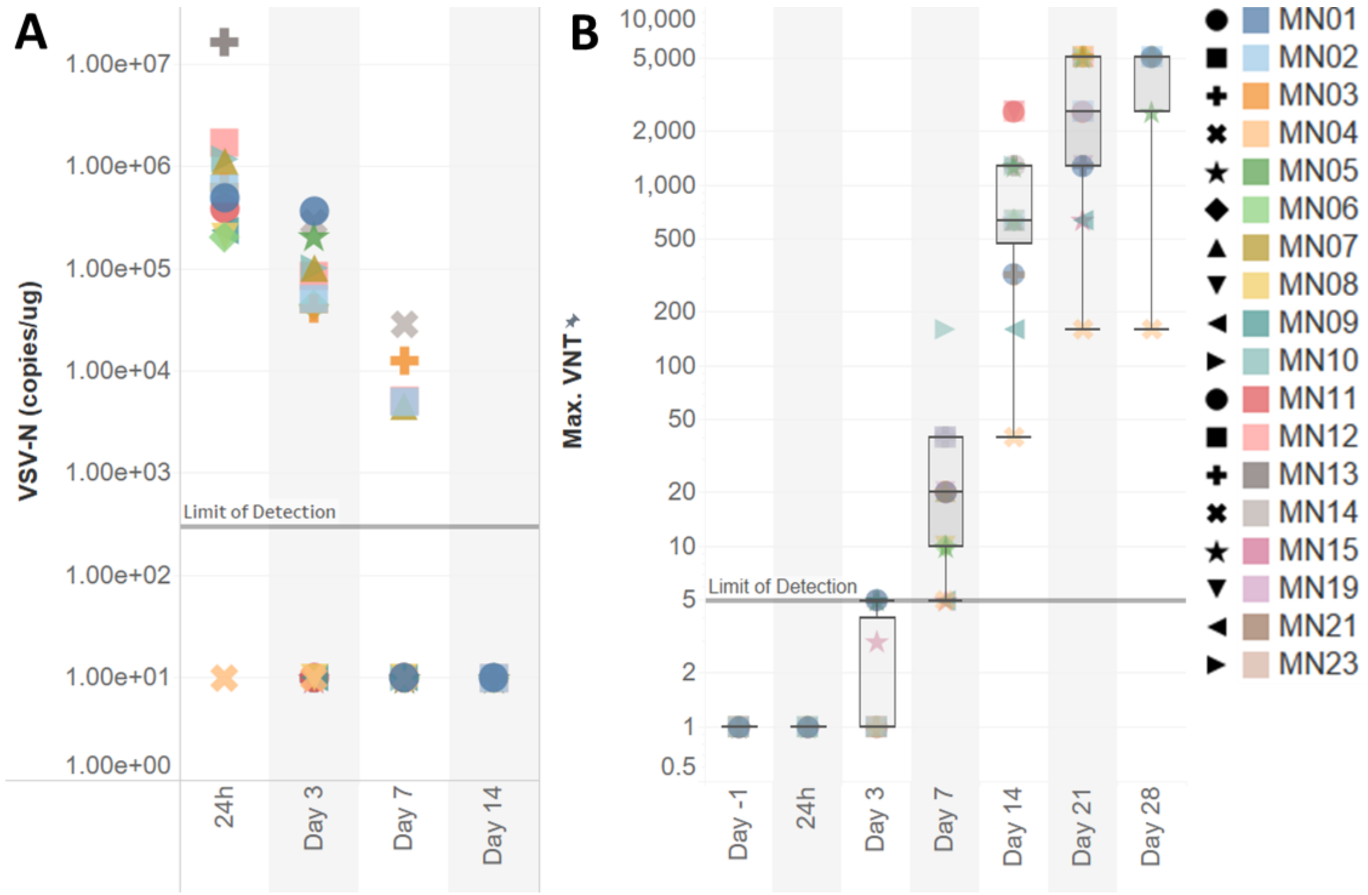

**Figure S6. Detection of virus shedding.** Detection of VSV-N RNA copies in RNA isolated from buccal swab, urine (separated into urine cells and supernatant), rectal swabs, and feces samples (from a subset of dogs).

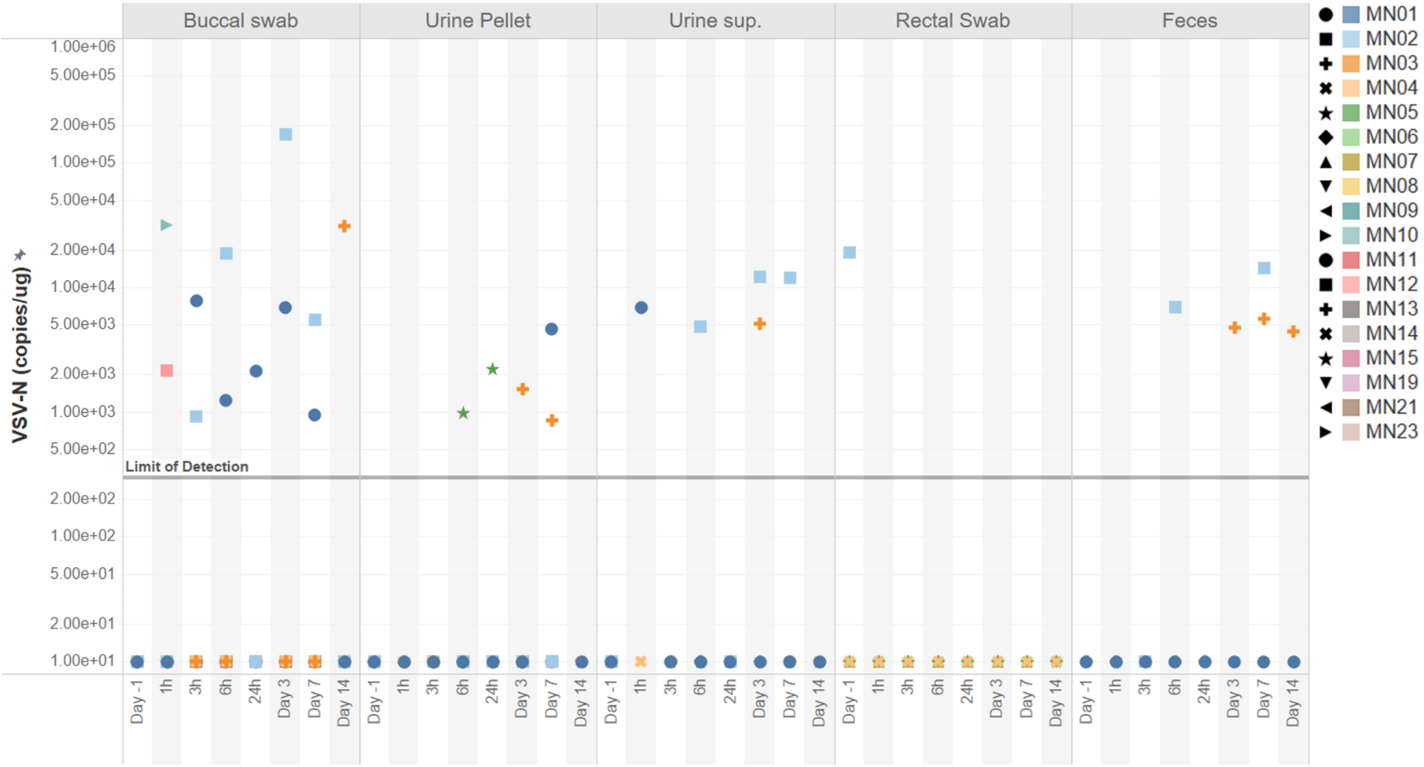

**Figure S7. Acute cytokine/chemokine responses after intravenous VSV treatment.**

Quantification of canine cytokines and chemokines (Granulocyte-macrophage colony-stimulating factor [GM-CSF], IL-6, monocyte chemoattractant protein-1 [MCP-1], keratinocyte-derived cytokine [KC], IL-2, IP-10, IL-8, TNF-α, IL-10, IL-7, IL-15, IL-18) in serum samples collected at baseline, 1, 3, and 6 hours post VSV infusion in the first 11 enrolled dogs.

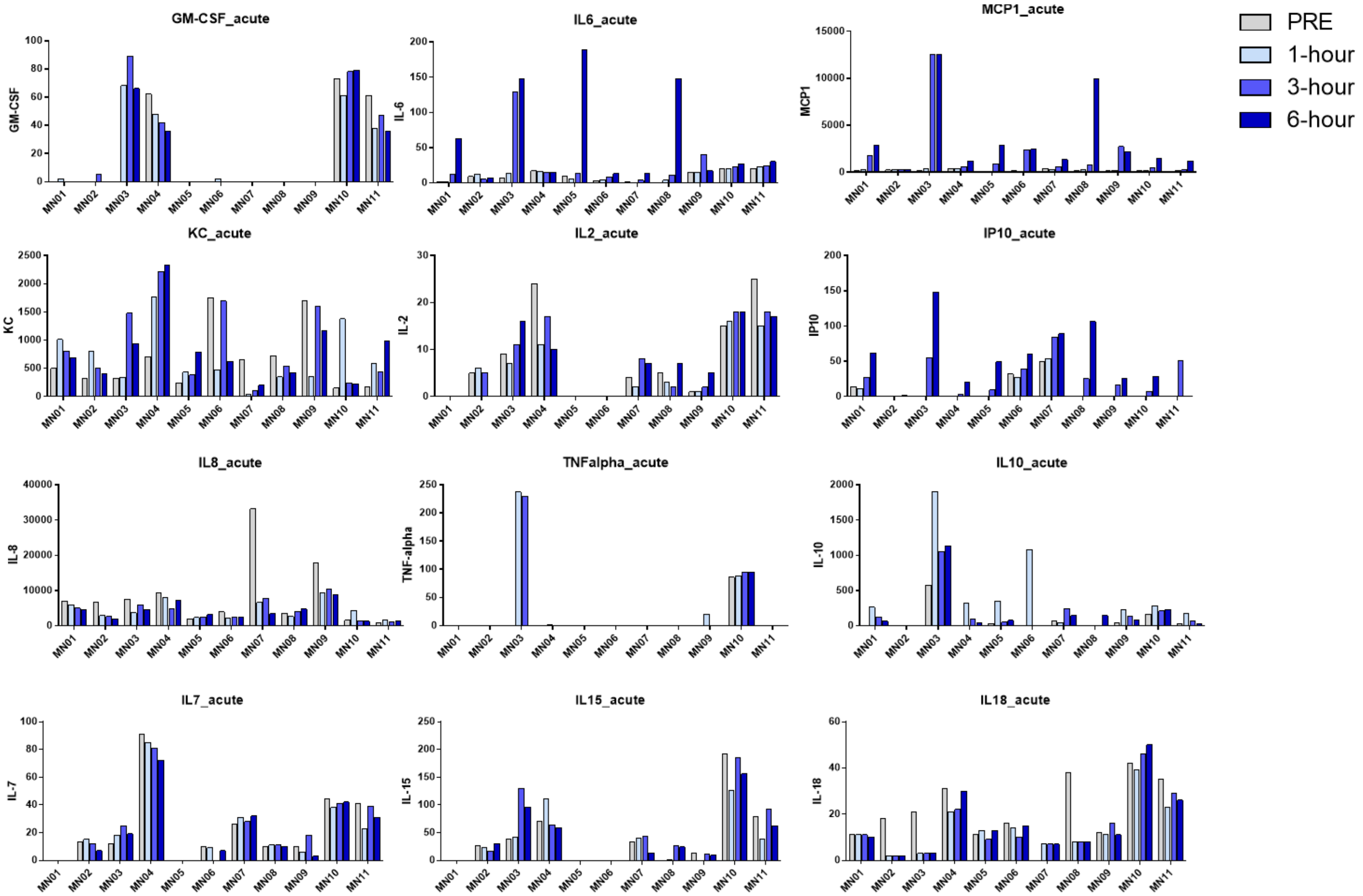

**Figure S8. RNAseq analysis of CCOGC control cohort.** (A) Unsupervised hierarchical clustering of RNAseq analysis data of available osteosarcoma tumor samples from the CCOGC control cohort. (B) Gene cluster expression summary score (GCESS) of osteosarcoma tumors. (C) Correlation of survival outcomes with CD37 monocyte (i), CD8 T-cell (ii), and cell cycle GCESS (iii), does not indicate a correlation between survival and CD8 T-cell GCESS in pre-treatment tumor biopsy samples (iv) in the CCOGC cohort.

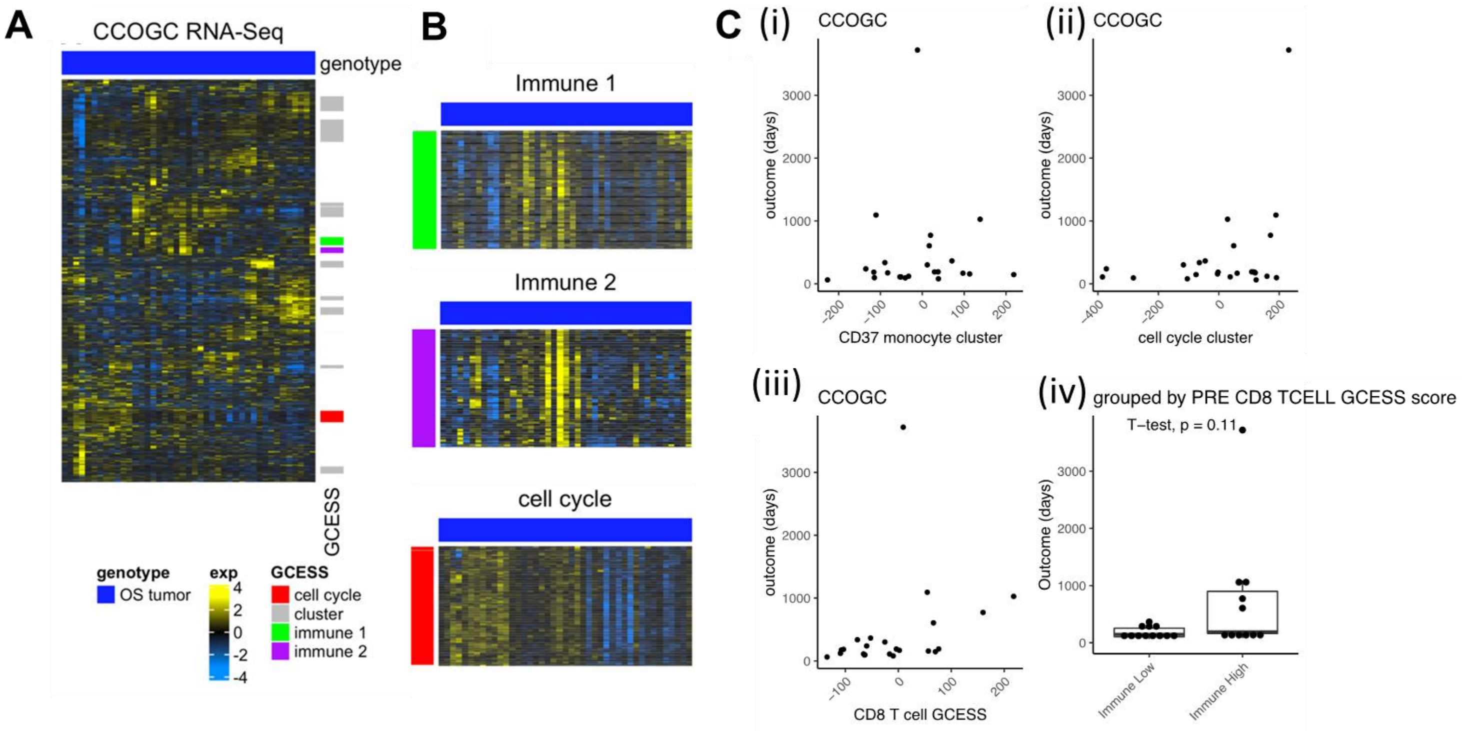

**Figure S9. Survival analyses comparing CCOGC and VIGOR cohorts.**

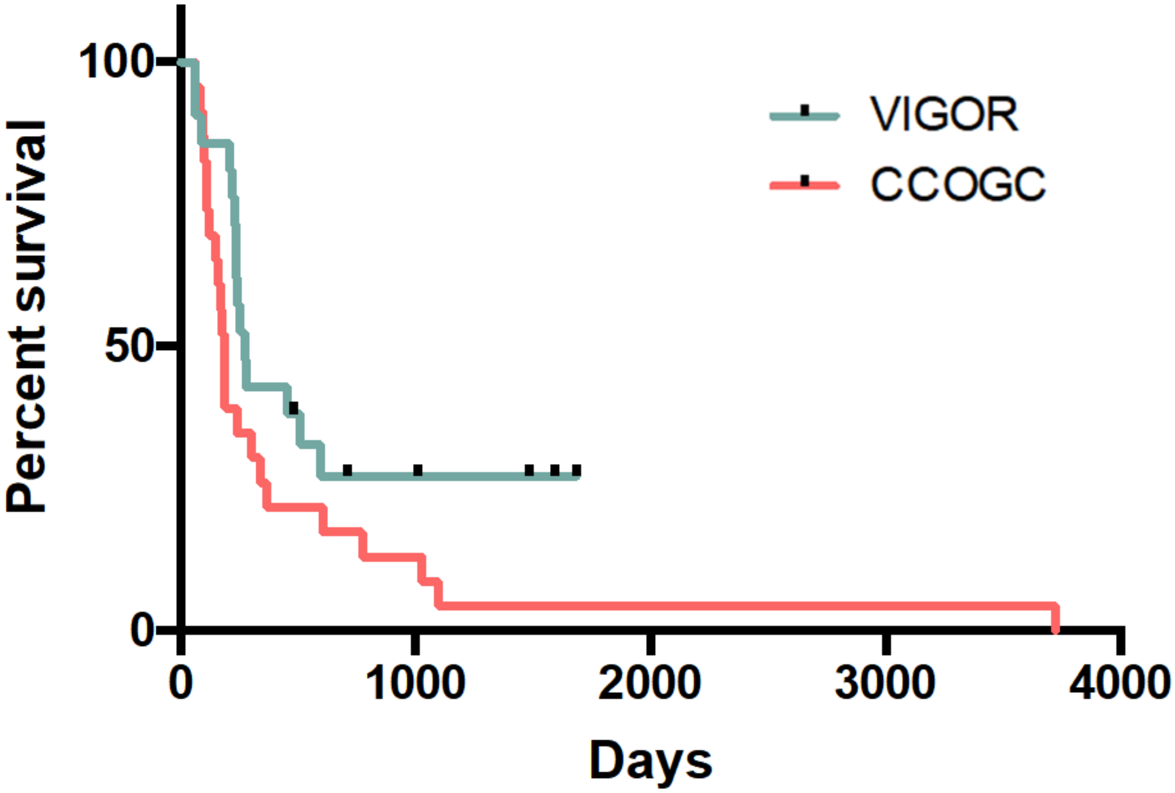
